# The Role of Cytoplasmic Interactions in the Collective Polarization of Tissues and its Interplay with Cellular Geometry

**DOI:** 10.1101/289520

**Authors:** Shahriar Shadkhoo, Madhav Mani

## Abstract

Planar cell polarity (PCP), the ability of a tissue to polarize coherently over multicellular length scales, provides the directional information that guides a multitude of developmental processes at cellular and tissue levels. While it is manifest that cells utilize both intra-cellular and intercellular mechanisms, how they couple together to produce the collective response remains an active area of investigation. Exploring a phenomenological reaction-diffusion model, we predict a crucial, and novel, role for cytoplasmic interactions in the large-scale correlations of cell polarities. We demonstrate that finite-range (i.e. nonlocal) cytoplasmic interactions are necessary and sufficient for the robust and long-range polarization of tissues — even in the absence of global cues — and are essential to the faithful detection of weak directional signals. Strikingly, our model recapitulates an observed influence of anisotropic tissue geometries on the orientation of polarity. In order to facilitate a conversation between theory and experiments, we compare five distinct classes of *in silico* mutants with experimental observations. Within this context, we propose quantitative measures that can guide the search for the participant molecular components, and the identification of their roles in the collective polarization of tissues.

## 1 Introduction

The development of a multicellular organism demonstrates the striking coupling of cellular states across large distances. While the study of gene expression has been the focus of research over the last 50 years, how these cellular states are coupled remains less explored, and at the forefront of research in developmental biology. As such, the coordination of cellular processes (e.g. cell division and rearrangements) on multicellular length scales are crucial to the emergent phenotype of an organism and requires the faithful transduction of directional information across tissues. Planar cell polarity (PCP) is understood to be the mechanism responsible for such tissue-wide signaling [1–8]. At the cellular level, polarity is defined as the asymmetric (i.e. anisotropic) localization of membrane associated proteins on the apicolateral cell junctions which is prompted and reinforced, in part, through cytoplasmic interactions and feedback loops [1, 3–6, 8–11]. Long-range polarization, then arises as a consequence juxtacrine signaling through which adjacent cells manage to align their polarities. In spite of the commonly accepted picture of the coupling between the cytoplasmic and intercellular interactions, the underlying mechanisms required for establishing the long-range planar polarity is yet to be elucidated [1, 3, 12, 13]. These intra- and intercellular interactions are largely carried out via two PCP pathways: “core-PCP” and “Ft/Ds” [3,6–8]. In order to construct our model, we adopt the core-PCP as a generic pathway, and use it as a reference system to clarify the phenomenology.

### Molecular components

The core-PCP pathway consists of six known proteins: Fz and Vang, which form complexes by binding to Fmi on the cell-cell junctions. Through formation of transmembrane Fmi-Fmi homodimers, the two complexes of Fz-Fmi and Fmi-Vang bind asymmetrically across the lateral junctions of two adjacent cells and form heterodimers [9, 11]. Additionally, within a given cell, Fz and Vang leverage the physical interaction between their respective cytoplasmic proteins (Dsh/Dgo for Fz, and Pk for Vang) to segregate and localize to the opposite sides of the cell [14, 15]. Although the presence of Dsh, Dgo, and Pk is found to be unnecessary for intercellular interactions, they facilitate segregation, and their absence impairs the long-range polarization [16–18]. One of the main goals of this paper is to address the significance of such cytoplasmic interactions in cellular polarization, as well as their interplay with juxtacrine signaling in establishing large-scale polarizations.

### Global Cues

Although the emergence of long-range polarization occurs through cell-cell interactions, external cues are believed to be necessary for fixing the direction of polarization [3, 4, 6, 18, 19]. The graded distribution of regulatory factors across a tissue, morphogens [6, 20, 21], mechanical signals [4, 5, 8, 13, 22–25], and geometric cues [26–30], are speculated to provide such global cues. Elongation in particular, has been observed to induce polarization either parallel or perpendicular to the axis of elongation. At a subcellular level, the polarization of microtubules and vesicle trafficking is also proposed to be acting as a bias to determine PCP orientation [29, 31–34]. In the mammalian cochlea and skin [19] polarization is perpendicular to the elongation axis. In mice, elongation along the medial-lateral axis has been suggested to orient the polarization along the anterior-posterior axis [30]. The present model aims to bring a mechanistic understanding to the potential role that cell geometry can play in the polarization of a patch of cells.

### Geometry and timescales

Several studies (like [35–37]) have proposed underlying physical mechanisms of PCP in ordered and isotropic systems. Establishment of long-range polarization during the course of development, however, can precede the formation of an ordered lattice, e.g. margin-oriented polarity in the prepupal *Drosophila* wing [5,12,24], suggesting that it is important to understand how PCP manages to propagate through disordered as well as elongated tissues. Quantitative measurements, in particular FRAP measurements of PCP proteins suggest turnover timescales to be much shorter than the timescales of cell re-arrangements **[ref: Personal communication Bellaiche]**. Thus, in order to study the characteristic patterns of PCP, it suffices to focus on a frozen geometry of tissue [5].

### Physical considerations

We find it crucial to clarify the term “long-range”, used frequently throughout this paper. The Mermin-Wagner theorem states that “true long-range” ordering is prohibited in 2D systems with continuous (e.g. rotational) symmetries, except at zero stochastic noise. The long-range order is referred to as the algebraic decay of correlation functions with distance. Below we will see that the magnitude of noise in our system drops as 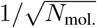, with *N*_mol._ the number of molecules participating in binding/unbinding reactions. Thus in the limit of large *N*_mol._→ ∞, the long-range order is achieved. For finite *N*_mol._, a state of quasi-long range order might exist.

### Outline and Results

The objectives of this paper are threefold. First, we describe a phenomenological reaction-diffusion model that includes the cytoplasmic interactions of like and unlike complexes, which couple a cell’s geometry to its polarity. We investigate the role of cytoplasmic interactions in establishing the long-range polarization, and show that nonlocal interactions mediated by diffusive cytoplasmic agents promote the segregation of unlike complexes and are crucial to stabilizing the global alignment of polarity. Furthermore we demonstrate that in elongated tissues, nonlocal interactions stabilize the polarization axis perpendicular to that of elongation. Finally, to demonstrate the significance of nonlocal interactions, and promote a conversation between theory and experiments, we study five types of *in silico* mutants within the context of the model and identify phenotypic similarities with observed mutants.

## 2 Model

We introduce a set of reaction-diffusion (RD) equations that describe the dynamics of heterodimer binding-unbinding at cell-cell junctions. Each cell is assumed to contain a finite pool of proteins A and B, which in their active state bind and form dimers at cell-cell junctions. The linear densities of total (bound plus free) A and B, are denoted by *a*_0_ and *b*_0_, and are assumed to be the same for all cells across the tissue. Assuming that transcriptional timescales far exceed the kinetic timescales of protein-protein interactions, *a*_0_ and *b*_0_ can be treated as time-independent.

At any point **r** on a junction shared by cells *i* and *j*, the concentrations of bound A and B localized on the *i* side are denoted by *u*_*ij*_(**r**) and *v*_*ij*_(**r**), respectively. Consistently with this notation, the corresponding concentrations on the *j* side are denoted by *u*_*ji*_(**r**) and *v*_*ji*_(**r**), such that *u*_*ij*_(**r**) = *v*_*ji*_(**r**). The key assumption in this model is that within each cell, the formation of a dimer at a point **r** is *nonlocally* enhanced by like dimers, and its dissociation is enhanced *nonlocally* by opposite dimers. In short, bound A nonlocally helps stabilizing of A and destabilizing of B; and vice versa. This represents an indirect positive feed-back between A and B in the adjacent cells: promoting (inhibiting) A in the same cell indirectly brings more (less) B to the other side of the junction. The nonlocal interactions are diffusively mediated through the cytoplasm; characterized by a single length-scale. The RD equations governing the binding/unbinding dynamics read:

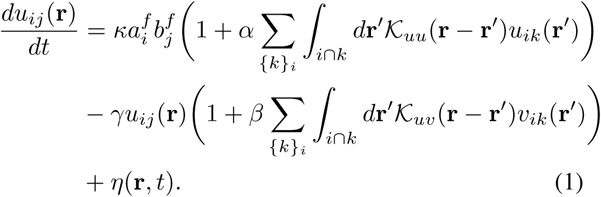

The notations adopted here are as follows: 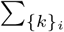 is the summation over all neighbors {*k*} of cell *i*; and ∫_*i*⋂*k*_ *dr*′ represents integration over the junction shared by cells *i, k*. In the above equation, the first and second terms on the r.h.s. correspond to the formation and dissociation rates, respectively. *κ* is the bare rate of formation. Based on the assumption that diffusion of unbound proteins is rapid we posit that the pool of free proteins are uniformly distributed on the perimeter of a cell. Therefore, the formation rate of *u*_*ij*_(**r**) is proportional to the densities of free A in cell *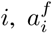*, as well as that of B in cell *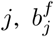*. In the second term, the dissociation rate is proportional to local concentration of the dimer itself, with the bare rate *γ*. The formation/dissociation processes are amplified by like/unlike dimers, respectively, through the nonlocal terms, *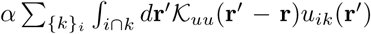*, and 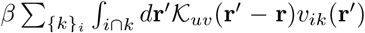 that characterize co-operative formation and dissociation. The functional form of the kernels 𝒦_*uu*_(**r**^*′*^− **r**) and 𝒦_*uv*_(**r**^*′*^ − **r**) and their coefficients *α* and *β*, are introduced below. Finally, the last term *η*(**r**, *t*) is a stochastic Gaussian white noise: ⟨*η*(**r**, *t*)⟩ = 00, and 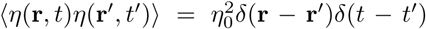, which arises from the molecular noise of chemical reactions and stochasticity in the upstream signaling pathways. The former, modeled as a Poisson process, is considered the dominant source of noise [36], with a magnitude scaling as 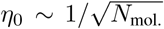 where *N*_mol._ is the number of molecules per area of the lateral interfaces. More precisely, the number of participating molecules is *N*_mol._ *ū/a*_0_, where *ū* is approximately the average value of *u*, and *a*_0_ = 1 is the unit of concentration. Using the variance of the number of reactions per unit time, that is given by the r.h.s. of Eq. (1), the noise level, is estimated to be of order *η*_0_ ≃ 0.01 – 0.1, for *N*_mol._ ≃1 – 5 × 10^3^, the approximate number of Frizzled molecules in the *Drosophila* wing [36].

The densities of free A and B are obtained by subtracting the densities of bound proteins from the total densities:

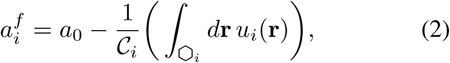

and a similar relation for *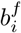*, by replacing *a*_0_ ↔ *b*_0_ and *u*_*i*_(**r**) ↔ *v*_*i*_(**r**). Here *u*_*i*_(**r**), *v*_*i*_(**r**) are the densities of dimers at point **r** on the boundary of cell *i*; *𝒞*_*i*_ is the perimeter of cell *i*. We define the cellular polarization with respect to the centroid of cell *i* at **R**_*i*_ (see Appendix (B)):

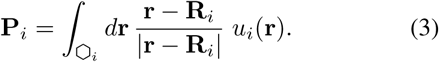

We calculate all the quantities in units where *a*_0_ = *γ* =ℓ = 1, where ℓ_0_ is the length of edges in an equilateral hexagon, or the average length of a potentially disordered cell. The coefficients *κ* = 10 and *α* = *β* = 5 are held fixed for all the cases discussed throughout the paper. In the following we will see that in certain situations, an alternative, but related, definition of polarization simplifies the analysis of the behavior of the system. We define two local variables: the cross-junctional polarity *p*_*ij*_(**r**) = *u*_*ij*_(**r**) − *u*_*ji*_(**r**) = *u*_*ij*_(**r**) − *v*_*ij*_(**r**), and the total concentration of dimers, *s*_*ij*_(**r**) = *u*_*ij*_(**r**) + *v*_*ij*_(**r**). Indeed, the second definition contains more information, and the cell polarities can readily be extracted from the junctional variables; see Appendix (B), for further discussion.

### Nonlocal Interactions

The kernels 𝒦_*uu*_(**r**^′^ − **r**) and 𝒦_*uv*_ (**r**_′_ − **r**), identify the functional form of the interactions between the concentrations of like and unlike dimers, respectively, and are taken to be exponentially decaying: 𝒦_*uu*_(**r**) = exp(−|**r**| */λ*_*uu*_) and 𝒦_*uv*_(**r**) = exp(−|**r**| */λ*_*uv*_), where *λ*_*uu*_, *λ*_*uv*_ are the characteristic length scales of *u*-*u* and *u*-*v* interactions, respectively. This functional form can be interpreted as an interaction mediated by diffusing cytoplasmic proteins with diffusion constant *D* andthe degradation rate *τ* ^−1^, such that *γτ* ≪ 1; thus *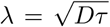*. The diffusion time of cytoplasmic proteins is of the order of state, as ∼10 min., much shorter than polarization dynamics which occurs on timescales of a few hours. We assume *λ*_*uu*_ and *λ*_*uv*_ are of the same orders of magnitude in the main text, and consider other cases in Appendix (D.4). The coefficients *α, β*, which parametrize the strength of cooperative interactions, are proportional to *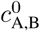*, the concentration of bound cytoplasmic protein to A and B at the junctions.

### Correlation Function

The correlation function of polarization is defined as a measure of alignment of polarity:

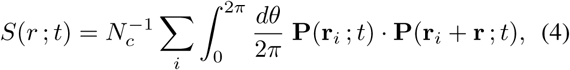

is the spatial average over the entire sample with *N*_*c*_ cells, at time *t*. Rotational invariance requires *S*(*r*; *t*) to depend only on *r* = |**r**|. The correlation length is thus defined as

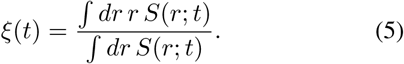

A simpler measure for the global *orientational* order is 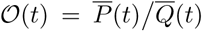, with *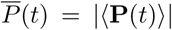*, and *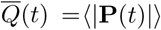*, in which ⟨•⟩ denotes spatial average. Thus 𝒪(*t*) saturates to unity for perfect alignment.

### Geometrical Disorder

The edge lengths are ℓ_*i*_ = ℓ_0_ + *ϵ*_*i*_, where *ϵ*_*i*_ *∈* [−*ϵ*_0_, + *ϵ*_0_] with *uniform* distribution, and 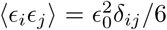; see Appendix (C). In 2D, lattice defects, i.e. non-hexagonal cells appear above a certain level of quenched disorder corresponding to *ϵ*_0_ ≃ 0.25. In dis-ordered cases, we use *ϵ*_0_ ≃ 0.5 and density of defects *n*_*d*_ *≃* 0.6, corresponding roughly to the statistics of larval and prepupal *Drosophila* wing [12]; see Appendix (C).

### Limit of Strictly Local Interactions (SLI)

In the limit of small *λ/*ℓ_*µ*_, we get for the kernels, *α*_*µ*_ *𝒦*_*µν*_ = 2 *αλδ*_*µν*_, where *δ*_*µν*_ is Kronecker delta. We define the coefficients of self-interaction, *α*_*s*_ ≡ *α*_*µ*_*𝒦*_*µµ*_ = 2 *α λ*, and similarly for *β*_*s*_, both of which are independent of *µ*. So in the SLI limit, the equations take the following form:

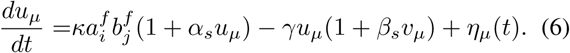

Before studying the real two-dimensional systems we briefly address the structure of steady-state solutions of the equation in 1D systems, in deterministic limit.

### 2.1 One-dimensional Arrays of Cells

Reviewing the one-dimensional case is valuable since it is more amenable to analytical treatment, hence some insight. Secondly, it captures some features of the 2D systems, in particular when the sixfold symmetry is broken, either explicitly by initial/boundary condition and/or geometrical anisotropy, or spontaneously, the system behaves much like a one-dimensional array with effective parameters. Therefore we first study the RD equations in one dimension in the mean-field (MF) approximation.

The ordered system at the MF level, i.e. *∀i* : ℓ_*i*_ = ℓ_0_, *u*_*i*_ = *u* and *v*_*i*_ = *v*, hence *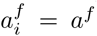* and *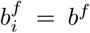*, exhibits a bifurcation from unpolarized to polarized the control parameter *b*_0_*/a*_0_, is increased above a critical value [38]. The MF polarization reads:

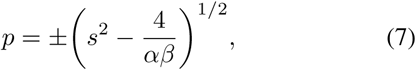

in which *s* = *u* + *v*. The bifurcation, thus, takes place at *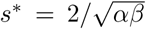* or in terms of actual control parameter: 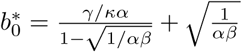. This result implies the divergence of the critical value *s*^*^ (or *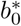*), for *αβ* → 0, indicating that the emergence of polarization requires cooperative interactions. Numerical solutions are presented in Fig. (2a), for a system with constant *A*_0_, *B*_0_, the total number of proteins per cell. Therefore, in a general disordered system with ℓ_*i*_ = ℓ_0_ + *ϵ*_*i*_, the concentrations *a*_0_ = *A*_0_*/*ℓ_*i*_ and *b*_0_ = *B*_0_*/*ℓ_*i*_ are randomized. Like a system with random critical point, the singularity is smeared out for *ϵ*_0_ ≠ 0.

**Figure 1:**
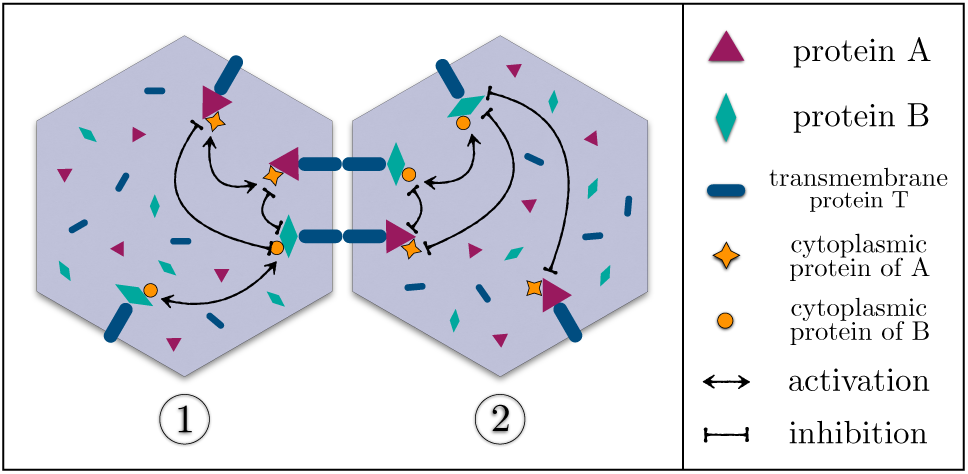
A schematic of the relevant cytoplasmic interactions. Membrane proteins A and B asymmetrically bind the transmem-brane proteins T, and form the dimers A-T : T-B across the junctions. At the junction shared by cells 1 and 2, two dimers of opposite directions are shown. The nonlocal interactions mediated by cytoplasmic proteins, couple the bound proteins A and B on different junctions. All pairs of complexes in a given cell, interact with one another, with exponentially decaying magnitudes; the like (unlike) complexes promote (inhibit) the membrane localization of one another. In order to keep the picture clear, we have not shown all the pairwise interactions, but only the generic ones.

**Figure 2:**
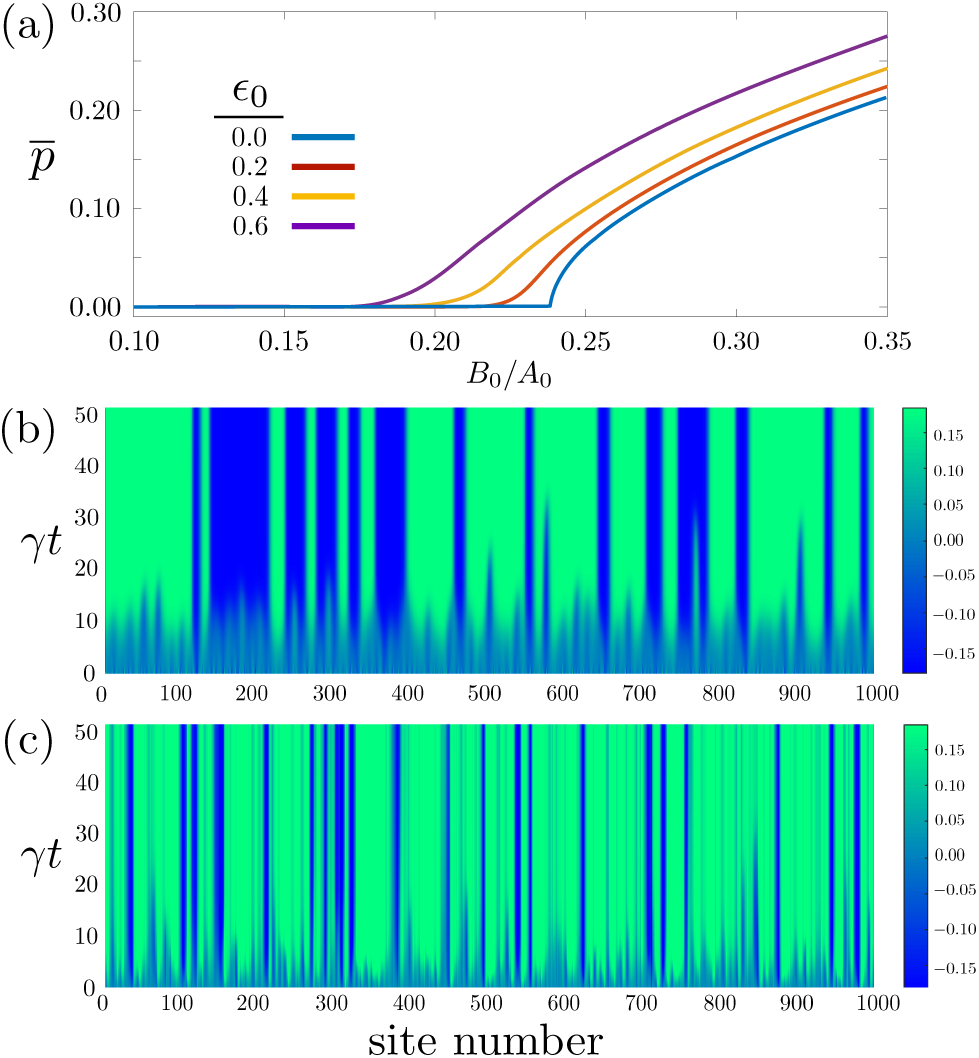
(a) shows numerical solutions of the average polarization against *B*_0_*/A*_0_, with *A*_0_, *B*_0_ the total number of proteins per cell, for different values of length disorder *ϵ*_0_ = 0 to 0.6. In ordered arrays, the critical value is *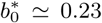*. The plot is obtained by ensemble averaging over 1000 realizations of quenched disorder, in arrays of 1000 cells. (b), (c) show the heatmaps of polarization of different sites in units of *a*_0_ℓ_0_, versus time (vertical axis) at *B*_0_*/A*_0_ = 0.3. (b) An ordered array with small bias, and (c) a highly disordered array *ϵ*_0_ = 0.6, with large initial bias.

From numerical solutions illustrated in Fig. (2b) and (2c), it is clear that in the limit of small stochastic noise and initial bias, the steady state is not guaranteed to be uniformly polarized in biologically relevant time scales of real systems. Initial imbalance of protein distributions is defined as *p*_0_ = |*u*_0_ − *v*_0_ |, with *u*_0_, *v*_0_, the spatial averages of initial dimers’ concentrations. The bias is defined as *δp*_0_*/p*_0_ ≃*σ*_0_, where *δp*_0_ is the magnitude of spatial fluctuations of initial polarity. Thus, small and large bias limits correspond to *σ*_0_ ≳1 and *σ*_0_ ≲1. While in ordered systems, a moderate initial bias suffices to achieve a uniform polarization, the patterns of polarity in highly disordered systems are robust and largely determined by the microscopic geometry of quenched disorder. Therefore we observe already in 1D, how quenched disorder imposes undesirable solutions, impairing the faithful transduction of information through PCP signaling. The situation gets only worse in two dimensions. One of the goals of this paper is to find mechanisms that circumvent this issue.

## 3 Two-dimensional Systems

The systems in one and two dimensions show inherently different behavior. In 1D, the proteins have only two junctions to localize at. This limited number of choices and the resultant predictability is absent in two dimensions. Due to the large number of possible steady states in 2D, the initial configuration influences the final state. We show in this section that within certain regimes of model parameters, NLI destabilizes a great portion of unpolarized fixed points, in favor of polarized ones. However, NLI comes with a drawback; increased sensitivity to the cellular geometry. Quenched disorder can thus have detrimental impacts on long-range polarity, since the individual cell polarity is coupled to local geometry. Therefore there exists a competition between the two effects: the benefit of segregation and disadvantage of sensitivity to local geometry. The latter is more pronounced in isotropic tissues, whereas elongation reduces this sensitivity. We find the optimal range of the NLI length scale, *λ*, that assists with establishing the long-range alignment.

### 3.1 Strictly Local Interactions (SLI)

The regime of SLI is defined as *λ/ℓ*_0_ →0, and corresponds to the short-range cytoplasmic interactions. In Sec. (2) we simplify Eq. (1) in the SLI limit. Here we present numerical solutions, but first we attempt to gain some insight using analytical approximate solutions. The definition of mean-field (MF) approximation in this system is uniform levels of *a*^*f*^ *b*^*f*^. This assumption is justified by the diffusive nature of *p, s* dynamics (see Ref. [38], and SI. Fig. (3a)). On the other hand, because of the symmetry of an equilateral system, where *α, β* coefficients are identical for all edges, we expect the steady-state *magnitudes* of *p, s* to be the same on all edges. Thus, three edges will be carrying inward and the other three carrying outward dipoles. The simplest of all MF solutions are those wherein translational invariance exists along each of the three axes separately, hence 2^3^ = 8 trivial configurations. The two types of trivial MF solutions can be seen in Figs. (3a1) and (3a2): sixfold symmetric configurations with nonzero net polarization, and two unpolarized states. There exist other types of solutions satisfying the MF criterion, in which translational invariance of polarity is not preserved; see Fig. (3b) for an example. We call these nontrivial MF solutions and are elaborated on in Appendix (D.3). It is easy to see that the nontrivial solutions, hugely outnumber the trivial ones (only 8 configurations). As such, starting from a random initial condition with no global cue, the system will tend to settle in a non-trivial fixed point, like the one depicted in Fig. (4a1). Since the analysis of nontrivial solutions provides intuitive arguments as to why SLI is insufficient to obtain long-range polarization, while NLI does indeed steer the system towards uniform polarization, we highly encourage the reader to peruse Appendix (D.3). Here, we only discuss the trivial solutions, since in the end we will see that NLI, suppresses a great deal of nontrivial solutions, as well as unpolarized trivial solutions. The RD equations in 2D are precisely the same as that in 1D, except the pool of proteins A and B is shared between six edges. For the polarized trivial MF solution, in isotropic hexagonal lattice, the net cellular polarization equals 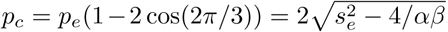, where *p*_*e*_, *s*_*e*_ are the values of *p, s* for one edge. For ordered lattices, the SLI critical point is thus *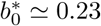*.

**Figure 3:**
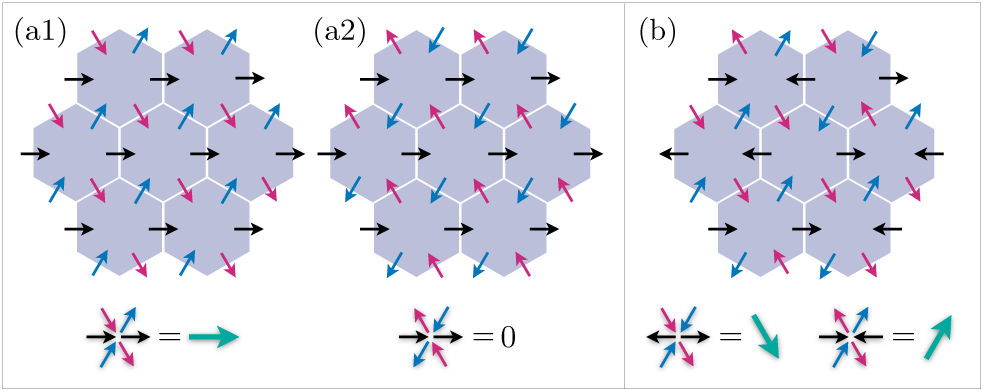
Cartoons of trivial (a1,a2), and nontrivial (b) mean-field solutions. In trivial solution, the translational invariance holds along each axis with nonzero (a1) and zero (a2) polarities. The latter is destabilized by any finite-range cytoplasmic interactions that induces segregation. In (b) the only constraint is uniform *a*^*f*^ *b*^*f*^ across the tissue, hence three incoming and three outgoing dipoles, and unequal cell polarities. In (b) the dipoles of the central cell and its right neighbor are shown for example.

**Figure 4:**
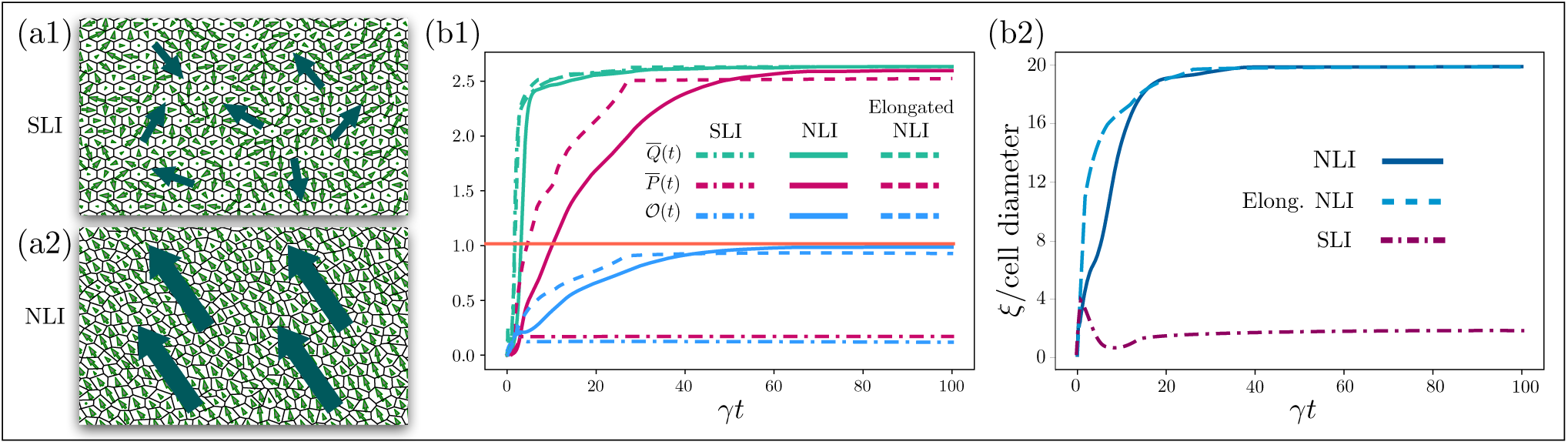
Generic steady states of two systems with identical parameters 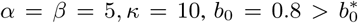 and random initial condition, for (a1) SLI limit (*λ* = *ℓ*_0_*/*100), and (a2) NLI of range *λ* = ℓ_0_*/*2, length disorder *ϵ*_0_ = 0.5, and *n*_*d*_ ≃ 0.6. The big arrows are to clarify the direction of dipoles. The time evolution of *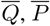*, and 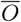, for isotropic systems with SLI and NLI, as well as elongated systems with NLI are shown in (b1). The curves corresponding to 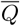 overlap to a large extent, which implies SLI systems are capable of polarizing individual cells, but fails to align the dipoles on larger scales. Comparing the curves of 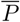, it is evident that alignment of cell dipoles requires NLI mechanism. This can also be seen in (b2) in which correlation length of SLI remains at around 4 cell diameters, whereas those of NLI systems grow until they reach the system size. The orange flat orange line in (b1) is the upper limit of 𝒪, which saturates at unity for perfect alignment. In all cases, the behaviors of all functions remains qualitatively the same for different realizations of quenched disorder and initial conditions.

Simulations of systems with random initial states and weak global cues, reveals that SLI is incapable of establishing the long-range alignment of polarity. A generic steady state of such systems, the time evolutions of *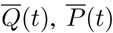*, and their ratio *𝒪* (*t*), as well as the correlation lengths *ξ*(*t*), are plotted in Figs. (4a1), (4b1) and (4b2), respectively. To simplify the comparisons, Figs. (4b1) and (4b2) include corresponding quantities of other cases of study, that are discussed in the next two sections. One important observation is the significantly <u>ra</u>pid dynamics of polarization of SLI case, in particular *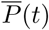*, acquiring its steady state value within a time scale of the order of ≲5(*γt*). This is due to the huge basin of attraction in SLI limit; any initial state is most likely very close to a fixed point, to which it is quickly attracted. This effect which happens at very small stochastic noise in SLI limit, is reminiscent of the glassy state, where quenched disorder provides a rough energy landscape making disordered configurations locally stable fixed points (more on this in the discussion section).

### 3.2 Nonlocal Interactions (NLI)

Nonlocal interactions, as introduced in Sec. (2), include finite-range interactions between the bound proteins within a cell. This interaction, in actuality, is mediated by cytoplasmic proteins, which enhance the segregation of membrane-bound proteins. For simplicity we take the form of interactions between like and unlike dimers to be identical. In one-dimension NLI leads to redefinition of *α*_*s*_, *β*_*s*_, due to self-interactions of edges and merely shifts the bifurcation point *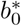*. The interesting role of NLI is revealed in two dimensions, among which are the alignment over large length scales, and the readout of geometrical information. Intuitively, the NLI facilitates the segregation of unlike proteins to two sides of the cell, by nonlocally attracting the like, and repelling the unlike dimers. Effectively, nonlocal interactions exclude the two MF solutions with zero-net polarizations in Fig. (3b2). Segregation makes the system behave more like a one dimensional lattice, by splitting each cell into two “effective edges”. For these reasons, the MF approximations with renormalized parameters, are more valid in systems with NLI compared to SLI.

While in relatively ordered tissues with NLI, and in the absence of an external cue, the orientation of polarization is determined purely by chance, namely stochastic noise and initial conditions, highly disordered systems show robustness against such random factors, and the fixed points of polarization fields are determined collectively by the geometry of the lattice. This stabilization of the fixed point becomes progressively more pronounced with increasing range of NLI and/or level of the quenched disorder. External cues of sufficiently large magnitudes can reorient the polarity towards the favored direction. Including a global cue in the form of constant gradient, we observe that NLI significantly enhances the susceptibility of the polarization field to the global cues. In particular, we find that, (i) the minimum strength of the cues required to rotate the polarity is much smaller in NLI systems compared to SLI, (ii) the response of the polarity is faster in NLI systems, and that (iii) unless the tissue is fairly ordered, *∊*_0_ ≲ 0.2, systems with SLI do not respond properly to the cues. As such NLI is a key to the detection of directional cues The impacts of the bulk and boundary cues are discussed in Appendix (D.4).

Using numerical simulations, with fixed aforementioned values of model parameters, we find a range of interaction length scale, 0.1 ≲ *λ/ℓ*_0_ ≲ 1, for which the NLI guarantees the alignment of polarization at large length scales. This range also depends on the distance from the critical point. We focus on the parameters deep in the polarized regime, *b*_0_*/a*_0_ = 0.8. A typical configuration of the steady states is illustrated in Fig. (4a2). The dynamics of *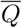* and *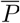*, shown in Fig. (4b1), implies that the emergence of collective polarization from an initially random cell dipoles, consists of two processes: (i) the segregation of PCP proteins within each cell and saturation of the amplitude of polarity, accompanied by the appearance of polarized domains of a few cells, which is followed by (ii) the subsequent coarsening and alignment of the domains on the tissue-wide scales.

Comparing the dynamics of SLI and NLI in Figs. (4b1) and (4b2), the role of cytoplasmic nonlocal interactions in cell-cell interactions, becomes immediately evident. Although polarization of individual cells is easily carried out in both SLI and NLI (see *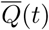*), long-range alignment of polarity (see 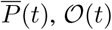 and *ξ*(*t*)) which requires intercellular communications, demands the presence of cytoplasmic interactions. Another important point to note is that the evolution of 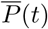 of systems with SLI and NLI, occur on remarkably different timescales. As briefly discussed in the previous section, in systems with SLI, the average polarization reaches its final negligible value within timescales ≲5(*γt*) corresponding to polarization of individual cells, because of the proximity of all initial conditions to a fixed point, and the immediate approach to the basin of attraction. On the contrary, the additional steps of coarsening and alignment of polarized domains in the NLI case, occur on timescales almost one order of magnitude longer than SLI, i.e. *≲* 20 – 50 (*γt*).

Finally, for *λ/ℓ*_0_ ≳ 1, all edges within a cell strongly couple to each other, thus the segregation cannot be accomplished, and the polarized state is unstable.

### 3.3 The Effect of Cell Elongation on Polarization

Elongation is considered, in many systems, a symmetry-breaking global cue. This is not due to a näive incorporation of length in the definition of the polarization, but the short junctions are indeed depleted of proteins [30]. Here we show that NLI endows the longer junctions with larger absorbing power than shorter ones, hence the stable perpendicular polarization.

The anisotropy of a cell i, is characterized by a traceless and symmetric nematic tensor, with the diagonal and off-diagonal elements, ±*ε*_*i*,1_ and *ε*_*i*,2_, respectively. The index of tissue elongation reads 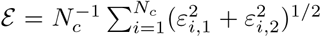; see Appendix (E.1). There are two possible choices for elongation axis. Elongation parallel to a pair of edges reduces the sixfold symmetry to a twofold associated to long junctions, and a fourfold, Fig. (5a); and vice versa if the elongation is perpendicular to a pair of edges; Figs. (5b,5c). In both cases, the system behaves qualitatively like a 1D problem extended in the direction perpendicular to elongation axis.

**Figure 5:**
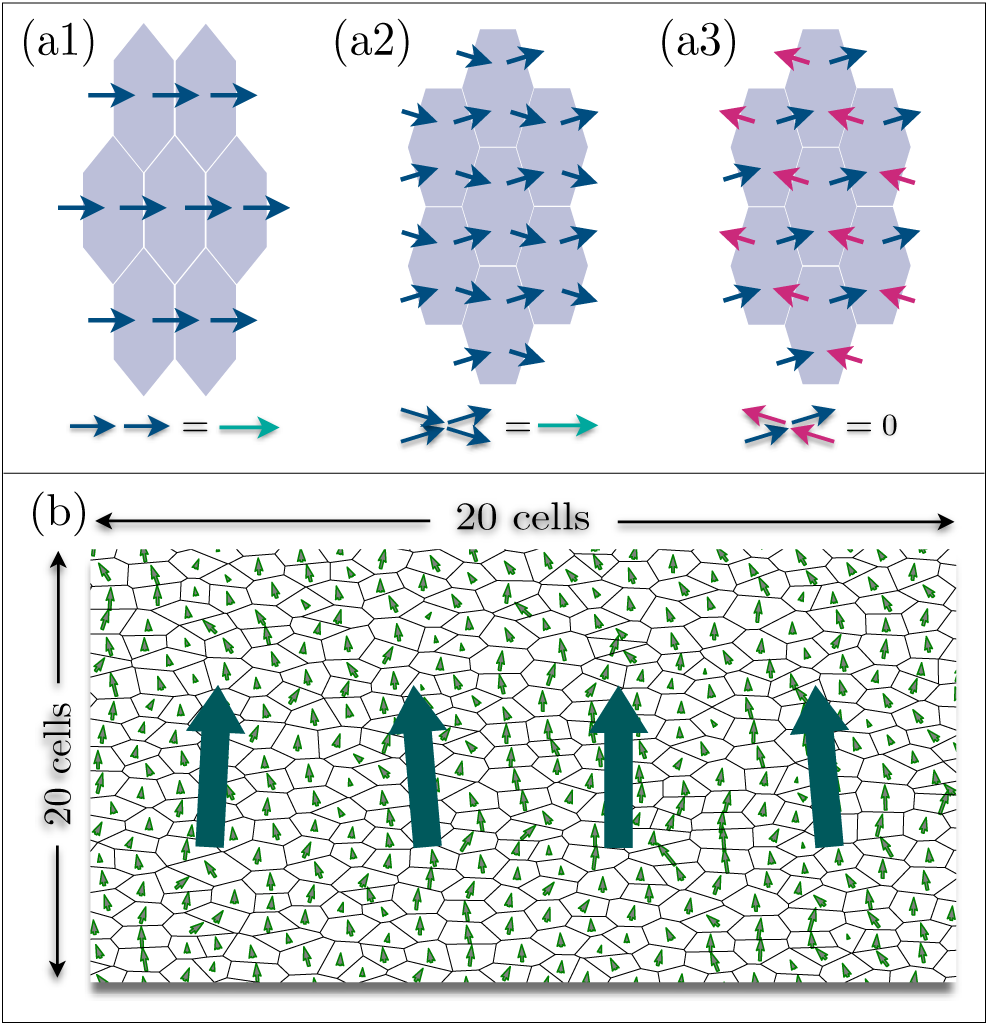
In (a1) the axis of elongation passes through a vertex, and the edges parallel to elongation wins the polarization competition; twofold symmetry like in 1D. (a2) and (a3) correspond to elongation perpendicular to an edge, therefore the two pairs of elongated edges compete: (a2) represents a polarized state, whereas (a3) is unpolarized. The latter is precluded by nonlocal interactions. (b) shows the final state of polarization in an elongated tissue along the horizontal axis, with *ε* = 0.4.

**Figure 6:**
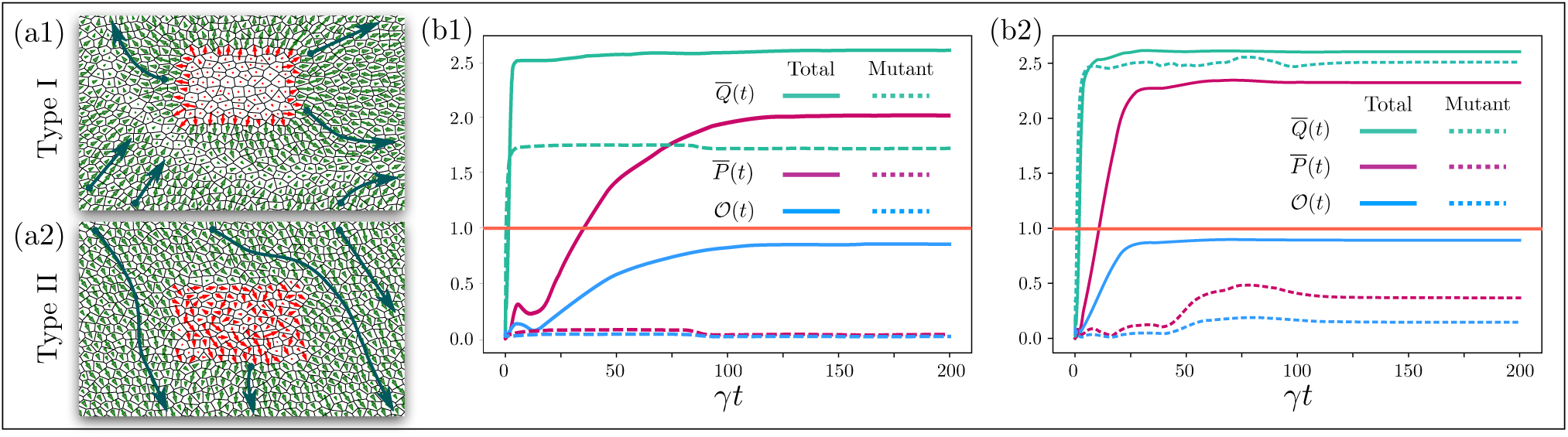
Steady-state of polarization in tissues with mutant clones at the center (red dipoles), depicted in (a1) type-I, with *b*_0_ = 0.1*a*_0_, and (a2) type-II with *λ/*ℓ_0_ = 0.01. (b1) and (b2) show the time evolutions of *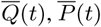*, and 𝒪 (*t*), for the whole tissue (solid curves), and mutant clones (dashed curves). Type I mutant patch lacks B protein, hence suppressed cell polarity. A proteins are attracted to the boundaries of the patch by B proteins in the surrounding WTs, and create radially outward polarization and non-autonomous formation of a crescent of vanishing dipoles. Type II lacks NLI; thus the cells are individually polarized like in SLI systems, but fail to align their dipoles. The WT dipoles avoid the mutant patch and become nearly tangent to the mutant-WT boundaries.

The elongated systems involve three different length scales and three regimes. With *L* the length of long junctions, we have: (i) *λ ≲ℓ*_0_ *< L*, (ii) ℓ_0_ *≲ λ < L* and (iii) ℓ_0_ *< L < λ*. For (i),(ii), NLI leads to the separation of positively and negatively polarized edges. For elongated cells, any subset of three adjacent edges includes one and only one long edge. From symmetry considerations, it follows that the longer edge is always the middle one of the three. Finally, the third regime *λ* ≳ ℓ_0_, *L*, is unstable, like the last regime discussed in isotropic case (*λ* ≳ ℓ_0_). Time evolutions of 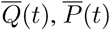 and 𝒪 (*t*) are shown in Fig. (4b1). We thus show that the observations of the experiments are consistent with nonlocal interactions. Furthermore the predicted onset of the perpendicular polarization as a function of elongation index, ε* ≃ 0.1, is in a very good agreement with that observed in the experiments [30]; see SI. Fig (4d).

### 3.4 Mutants and The Associated Phenotypes

PCP mutants exhibit lack of orientational order of e.g. hairs, bristles. that are induced either autonomously or non-autonomously by the mutant clone [6, 10, 39–41]. The phenotypes are commonly used to specify the role of the corresponding protein in the PCP pathway. Here we introduce possible classes of mutations within the framework of our model, and explore their phenotypes. In our model the random orientation or lack of polarization can be the signature of, (I) lack of sufficient protein B, i.e. *b*_0_*/a*_0_ ≪1; and/or absence of NLI, i.e. *λ/*ℓ_0_ ≪1. In the main text, we investigate the properties of these two mutant types, and ask whether or not shortage of a membrane protein, or the suppression of NLI in a patch of cells, reproduces the observed phenotypes. The results of numerical solutions are illustrated in Fig. (4a1) and Fig. (4a2). In Appendix (F), we discuss three more mutant types. Type III corresponds to lacking cytoplasmic proteins; type IV, to irregular and severed cell geometries; and type V, to double-mutant lacking both membrane proteins. Type IV is of great interest as it was suggested in [24], that in the absence of global cues, e.g. *ft*^−^ in *Drosophila* wing, irregular cell packing disrupts the polarity. Experiments on double mutants *Vang*^−^*fz*^−^, suggest that they show less non-autonomous phenotypes compared to single mutants [16,42]. Thus within the framework of our model, we examine double-mutants, in which both membrane proteins A and B are lacking, and verify that the prediction of our model is consistent with experiments. The comparison of double and single mutants are crucial to our understanding of the mechanisms of the intercellular signaling [18]. We present and discuss type-I, II and III mutants in Appendix (F.1), (F.2) and (F.3), respectively. Finally, under certain circumstances, topological defects and domain walls appear in tissues, either as transient or permanent polarity defects. These effects and their corresponding figures are discussed in Appendix (F.4).

The two types of mutants I and II, can be discerned both qualitatively and quantitatively. In Figs. (6b1) and (6b2), we see that in both cases, the mutant clones have small net magnetization compared to the surrounding WT cells: *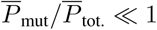*. In type I, this is due to inability of individual cells to polarize due to lack of B proteins, whereas in type II is a consequence of ra<u>n</u>dom orientation of dipoles. The average of magnitudes *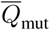*, behaves differently in type I and II mutants. In type I, the magnitude of dipoles in the bulk of mutant patch is distinguishably smaller than WTs 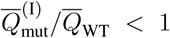, thus the hairs grow rather apicobasally with no preferred orientation; Fig. (4b1). The largest contribution to ^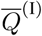^ comes from the mutant boundary cells, with radially outward dipoles. This is checked by enlarging the size of the patch such that the ratio of the number of cells at the boundary to that of the bulk of the patch drops. In type-II mutants, the polarizations of individual cells are compa<u>ra</u>ble to those of WTs, but are in random directions; like *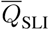* in Fig. (4b2). In type-I, the mutant cells not only fail to polarize, but also distort, non-autonomusly, the WT polarization at a distance of a few cell diameters from the mutant boundary. The ring of emanating dipoles at the clone’s boundary arises from the localized A proteins, attracted by the surrounding WTs (note that polarization is defined in terms of localized A, hence outward dipoles). Unlike type I, the non-autonomous effects of type II are minimal, causing minor deflections of dipoles enclosing the mutant patch, with WT dipoles’ orientations more or less tangent to the WT-mutant borderline. Interpreting the two mutant types, our model predicts non-autonomous phenotypes in the absence of membrane proteins (type-I), while lacking cytoplasmic diffusive interactions causes only autonomous phenotypes. These results are in good agreement with the experimental observations of *Fz*^−^ (or *Vang*^−^), and *Dsh*^−^, i.e. the membrane and loss of function of cytoplasmic proteins in the core-PCP, which show non-autonomous and autonomous phenotypes, respectively [6, 9, 16, 24, 35, 39, 42, 43].

We believe that our analysis of mutant phenotypes can be used as a reliable diagnostic for characterization of the role of various PCP components. The significant differences in non-autonomous effects on the polarization patterns of the WTs, in particular the radial orientation of dipoles in type I, and the crescent of vanishing polarization in the WT region is a fast qualitative way of identifying the source of mutation. Furthermore, the ratio *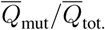*. can serve as a qua<u>n</u>titative indicator of mutant types. What observable does *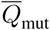* represent in an experiment? In unpolarized cells, hairs grow perpendicularly to the apical surface (i.e. along *z*-axis). The in-plane projection of the hair lengths that is easily observable in all images, is thus a measure of the average magnitude of polarization *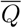*, and can be used in the taxonomy of mutants, and identification of the role of the corresponding component in the PCP pathway.

## 4 Summary and Discussion

In an attempt to understand the connection between cell-cell communications and intracellular interactions, and based upon a few realistic assumptions deduced from the recent experimental studies, we devised a generalized reaction-diffusion model by incorporating the nonlocal (i.e. finite-range) cytoplasmic interactions, into the interactions between membrane proteins. In line with our phenomenological approach we remain agnostic to the details of the cell biological processes, but we suggest that cytoplasmic proteins such as Dsh, Dgo, and Pk, could potentially mediate these interactions. Although the functional form of interactions are assumed to be exponentially decaying (diffusion of a degrading protein), any finite-range kernel would produce qualitatively the same results. Assuming *λ*_*uu*_ = *λ*_*uv*_, and for the specific set of model parameters we chose, the optimal range of interactions to achieve long-range polarization is found: 0.1 ≲ *λ/*ℓ_0_ ≲ 1. In Appendix (D.4), we demonstrate the results of the simulations for cases where the ranges of interactions *λ*_*uu*_ and *λ*_*uv*_ are incomparable. Although in this paper we defined and used dipole-dipole (including the magnitudes) correlation functions, the angular correlations show very similar behavior (results not shown). We further examined the response of polarization to external cues, and concluded that, NLI systems are more efficient in detecting and responding to the weak global cues, and indeed, are essential to the orientational order of the collective polarity of the tissues, that are moderately disordered, e.g. before the formation of the hexagonal order.

Since the PCP organization occurs in two dimensions, we find it crucial to make a few clarifying comments on the relevance of the Mermin-Wagner theorem to our system. Firstly, we emphasize that the lack of order in SLI limit is fundamentally outside the scope of Mermin-Wagner theorem, in which the entropic cost of the long wavelength fluctuations prohibits “true” long-range ordering, i.e. algebraic decay of correlation length, in 2D systems with continuous symmetries. For fixed other parameters, the phase diagram of our model consists of two control parameters, (i) *η*_0_, and (ii) *λ/*ℓ_0_. Whereas in the NLI regime, there exists a transition from disordered to ordered phase as the temperature is lowered (not explored here), the SLI limit does not show ordering even at zero temperature, i.e. strictly deterministic dynamics. As was shown in Fig. (4b1), in both cases of SLI and NLI, the first stage of dynamics corresponds to saturation of the spin magnitudes, after which the system behaves (qualitatively) like a 2D ferromagnet. In analogy with the XY-model, the spin-spin coupling constant in our system is an increasing function of *λ/*ℓ_0_, which is negligible in the SLI limit. Furthermore, it is noteworthy that unlike spin glasses, the glass-like state of the SLI limit is not induced by the quenched (geometrical) disorder. Indeed, SLI’s glassy state exists even in perfectly ordered systems.

We next investigated the role of cell elongation as a symmetry-breaking cue, and showed that NLI endows the longer junctions with more “protein-absorbing” power, and that this effect is beyond the superficial shape effect arising from the definition of polarity. Intuitively, the elongation drives the behavior and the fixed point of the system towards an effectively 1D array of cells. We shall emphasize that this prediction is only valid under the following assumptions: (a) polarization is predominantly induced by reaction-diffusion processes, and (b) no mechanical dynamics (like tissue flow or cell division) are involved. In other systems, e.g. wing of *Drosophila*, where the polarization is observed to be parallel to the elongation axis the polarization of microtubules is believed to be the dominant mechanism [33]. The relative rates of cell division and PCP relaxation, is another parameter that can induce perpendicular or parallel polarizations [5].

Finally we examined the predictions of our model in the cases of five mutant types, in which (I) one of the membrane proteins (B) is lacking, and (II) the nonlocal cytoplasmic interactions are absent, hence a clone with SLI, (III) the cooperative interactions are suppressed, (IV) geometrical irregularity is enhanced, and (V) both membrane proteins (A and B) are missing, i.e. double-mutants. The corresponding phenotypes were identified, and for distinguishing type I from II, we proposed a measure, i.e. average in-plane projection of the hair length. The results, as discussed in Sec. (6) exhibit good similarity with experimental observations, and lend more support to the significant role of NLI in the signaling pathway. In Appendix (F), we discuss the other three types. Type-III in which cooperative interactions are suppressed is interpreted as mutants lacking of cytoplasmic proteins. It is important to distinguish type II from III, both of which involve cytoplasmic proteins. Type II corresponds to loss of function (diffusion) of proteins like Dsh that mediate the nonlocal interactions, yet the local interactions are not impaired. In type III, the cytoplasmic proteins are absent altogether. Type IV is also of great interest, as it was suggested in [24], that cell packing impairs Fz feedback loops in the absence of global cues, i.e. Fat, in the *Drosophila* wing. Finally, the double mutants lacking both membrane proteins A and B are investigated in type V. Interestingly, the type-V phenotypes show less non-autonomy than single mutants (type I), which is consistent with the observations in experiments. The results of all simulations are presented in Appendix (F).

Although our model is not the first phenomenological approach to the problem of PCP, we claim that the features included in this model, and its predictions, capture a broader range of recently observed phenomena. Some of the successful previous studies (e.g. [18, 38]) consider one-dimensional arrays of cells. Although the general frameworks proposed in such studies have provided us with valuable insights, two-dimensional systems have to overcome the issues associated with the large number of fixed points in SLI. Other successful models such as that studied in Ref. [13], infer effective interactions from the observed response of the polarization to a combination of processes; cell elongation, cell rearrangements, and divisions. However the model does not aim at explaining the underlying mechanisms of emerging polarity at the cellular level. Among the models derived from subcellular interactions, the one put forward by Burak and Shraiman [36], is closely related to ours. In spite of the similarities in the general mathematical approaches to the problem, there exist differences at the level of phenomenology of interactions, e.g. in the direct vs. indirect implementation, as well as the locality/nonlocality of the cytoplasmic interactions. Furthermore, having included the nonlocal cytoplasmic interactions we found it crucial to address the impacts of geometrical information such as quenched disorder and tissue anisotropy. Finally, we studied the possible mutants within the scope of our model. Former studies, such as in Refs. [24] and [35], have investigated the phenotypes induced by irregular geometry as well as domineering nonautonomy, and proposed mathematical models with parameters inferred from measurements. Adopting a phenomenological approach, we tried to keep the number of model parameters as few as possible in order to determine the minimal set of criteria to achieve the large-scale PCP alignment. The successful reproduction of the perpendicular axes of polarity and elongation, as well as some of the experimentally observed phenotypes, suggests that our phenomenological model captures the salient features of the generic cytoplasmic and intercellular interactions, and has the predictive power of identifying the roles of various proteins in PCP pathways.

We believe that, in spite of several attempts to understand the mutual interplay of PCP and tissue mechanics, a unified perspective of the relevant physical mechanisms is yet to be discovered. Therefore, the mutual signaling of mechanical properties and polarity, both of which depend on the geometry and contribute to the dynamics of the tissue flow is among the most important problems to be considered in the future. Our study lays the groundwork for further investigations in this direction, by clarifying the response of polarization to the cell geometry of the tissue.

## ACKNOWLEDGEMENTS

The authors are grateful to B.I. Shraiman for helpful discussions. S.S. was supported by the Gordon and Betty Moore Foundation through Grant GBMF2919.01. M.M. would like to thank the Simons Foundation MMLS program for support.

## Appendix

### A The Model and Its Ingredients

We begin by writing the full reaction-diffusion equations for the binding/unbinding dynamics of the local density of the complexes complexes [A:B], namely *u*_*ij*_ (**r**). The corresponding equation for [B:A] or *v*_*ij*_ (**r**), is obtained by simultaneous replacements: *a* ↔ *b* and *u* ↔ *v*.

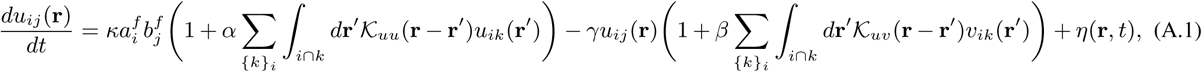

where *γ*^−1^ is the timescale associated with complex dissociation. As mentioned in the main text, the kernels 𝒦 (**r**) are assumed to be of the form of exp(−|**r**| */λ*), which was motivated by the diffusive nature of the cytoplasmic proteins carrying the interactions. Although the kernels couple the concentrations of the complexes on the boundaries of the cells, the coordinate **r** can in principle represent any point within the cytoplasm as well as the junctions. Suppose that a cytoplasmic protein C, obeys a diffusion equation with degradation time *τ* :

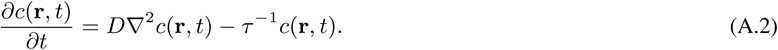

Assuming the degradation and hence dynamics of C takes place on a much faster timescale than that of PCP, i.e. *γτ*≪ 1, it suffices to only consider the steady-state solutions of protein C. Now for C_A/B_ cytoplasmic protein of A/B we have (A/B means A or B):

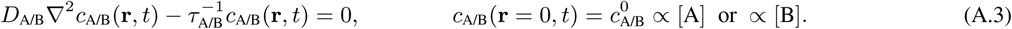

Here, **r** = 0 corresponds to the specific junction on which the concentration of A is measured, and from which C diffuses. Superimposing the concentrations of C at a given point **r**, the total amount of C emanating from all points around the cell reads:

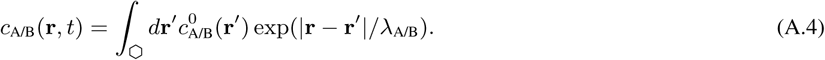

Here, *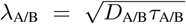* are the diffusion length of proteins C_A_ and C_B_. The diffusing proteins enhance the formation of like complexes and suppress that of unlike complexes. This nonlinear effect in turn, depends on the respective interactions with the target complexes. Altogether, the coefficients are lumped into the phenomenological constants *α* and *β*.

For notational convenience, in the following paragraphs greek letters label edges, e.g. *µ* ≡*i* ⋂ *j*. Using Eqs. (A.1) and the definitions of *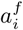* and *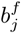*, we can solve for the dynamics and the steady states of *u*_*µ*_(**r**). Evidently, solving the above integro-differential equations is a cumbersome task. One possible simplification is “junctional” averaging: *u*_*µ*_ = ∫ _*µ*_ *d***r**′ *u*_*µ*_(**r**′)*/*ℓ_*µ*_, in terms of which we have:

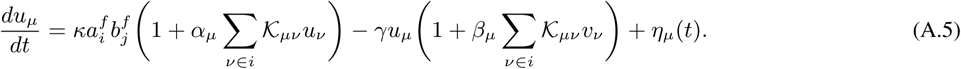

In the above equation, *α*_*µ*_ = *α/*ℓ_*µ*_, *β*_*µ*_ = *β/*ℓ_*µ*_, and for the kernels *𝒦*_*µν*_ = *𝒦*_*νµ*_ = ∬_*µ,ν*_ *d***r***d***r**^*′*^ *𝒦* (**r** *–* **r**^*′*^). The diagonal elements equal

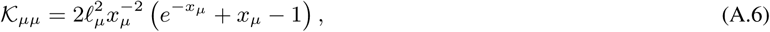

in which *x*_*µ*_ = ℓ_*µ*_*/λ*. Finally, the stochastic noise is renormalized: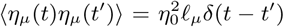. We see from the above equation and Eq. (A.1) that all the points on a given edge satisfy the same equations, and are only distinguished from points on other edges by the term *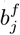*, i.e. the amount of free B in a neighboring cell *j*. Therefore, in this model, the assumption of uniform density on a junction is a reasonable one, provided *λ/*ℓ is small enough that different junctions do not interact. It is noteworthy, however, that in reality the core proteins for example Flamingo and Frizzled in the prepupal and pupal wing of *Drosophila*, are observed to be persistently localized at subdomains of plasma membranes, called “puncta” [44].

#### Limit of Strictly Local Interactions (SLI)

In the limit of small *λ/*ℓ_*µ*_, we get for the kernels, *α*_*µ*_ *𝒦*_*µν*_ = 2 *α λδ*_*µν*_, where *δ*_*µν*_ is Kronecker delta. We define the coefficients of self-interaction, *α*_*s*_ ≡ *α*_*µ*_ *𝒦* _*µµ*_ = 2 *α λ*, and similarly for *β*_*s*_, both of which are independent of *µ*. So in the SLI limit, the equations take the following form:

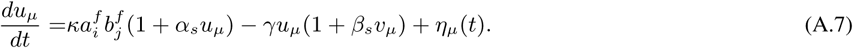

Before discussing the mean-field solutions we introduce the precise definitions of polarizations and their relations.

### B Definitions of Junctional and Cellular Polarity

Planar cell polarity can be defined at either junctional or cellular level. For individual junctions, polarity is defined as the difference between the concentrations of *u*_*ij*_ = [A_*i*_ : B_*j*_] and the opposite dimer, *u*_*ji*_, thus *p*_*ij*_ = *u*_*ij* −_ *u*_*ji*_ = *u*_*ij*_ − *vij*. This is indeed what we use in the 1D case in main text. This quantity, along with the sum *s*_*ij*_ = *u*_*ij*_ + *uji* = *u*_*ij*_ + *vij* of the concentrations of the dimers, contain the same exact information as *u*_*ij*_ ‘s. In other words, given the concentrations of dimers, one can calculate the junctional polarization and sums, and vice versa. Below we see that cellular polarity is obtained by integrating the concentration of A_*i*_:B_*j*_, i.e. *u*_*ij*_. Thus, the cell polarity can be equivalently computed using the junctional dipoles and sums.

Cellular polarization is referred to as the asymmetric (anisotropic) distribution of PCP proteins in the cells. The precise mathematical definition of cellular polarization can be ambiguous, due to the lack of knowledge about the exact subcellular mechanisms through which a cell gathers information from proteins on its periphery, and determines the direction of e.g. hairs or bristles. However, for phenomenological purposes, all definitions capture the features of interest. In spite of this freedom in defining the cell polarity, we realize that after all, polarization is a cellular property. Cell polarity, depending on the symmetry of binding complexes, might be either a vector quantity, called vectorial polarity, or a nematic, which is then called axial polarity. Vectorial and axial polarities are used when PCP proteins form heterodimers and homodimers, respectively. The former is a vector identified by a magnitude and angle ∈ [0, 2*π*), whereas the latter is a traceless nematic tensor with a magnitude and an angle ∈ [0, *π*). In our case of study, the polarization involves two distinct complexes, A and B, hence vectorial PCP. However, we shall introduce both vectorial and axial cell polarities.

#### (i) Vectorial Polarization

This definition is used to calculate the “dipole moments” of the distribution of bound A (or B), around each cell. The polarization vector associated with cell *i* is defined as:

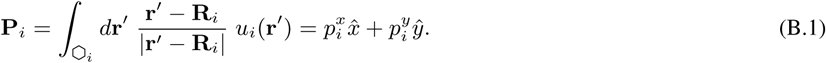

Here **R**_*i*_ is a reference point within cell *i*, with respect to which the polarization is defined. Since the total amount of bound proteins is nonzero, the dipole moment depends on the reference point; we take this to be the geometrical center of mass of each cell. One can alternatively define polarizations in terms of distribution of B proteins, i.e. *v* dimers. The polarization vector is defined by a magnitude and and angle *θ*_*i*_ ∈ [0, 2*π*) from *x*-axis:

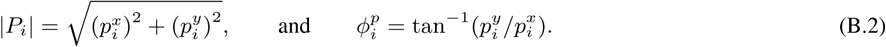

The polarization of the tissue with *N*_*c*_ cells, and its global order are characterized by the following quantities: (1) average polarization,

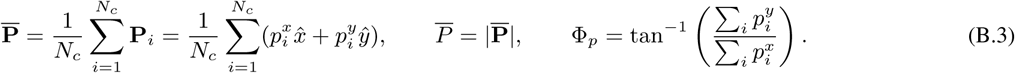

(2) average magnitude of polarization, and (3) the ratio of (1) and (2),

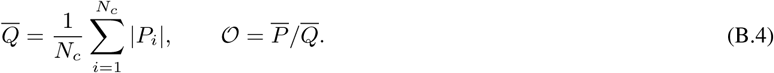

The ratio *𝒪* approaches one when the system is perfectly aligned.

#### (ii) Axial Polarization

The second definition is a measure for cellular polarity that is used especially when dealing with axial nematic PCP (like Celsr (or FmI) homodimers) which merely determines the axis of polarization by measuring the traceless nematic tensor of the polarity:

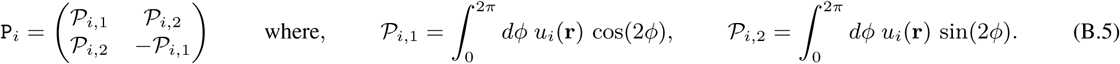

In the above equation, *ϕ* is the polar angle of point **r** (with respect to the centroid), on the periphery of cell *i* and is measured from a reference axis. The concentration of bound A at point **r** is denoted by *u*_*i*_(**r**). In using these formulae for inferring the polarization from experimental data, *u*_*i*_(**r**) is replaced by the intensity of light reflected from a GFP at point **r**. The magnitude of polarization equals *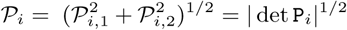*. Its orientation is determined by angle *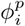* satisfying 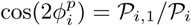 and 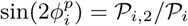. In Appendix Sec. (E.1), we use the same general formalism for the cell shape nematic tensor.

### C Measure of Quenched Disorder and Topological Defects in 2D Systems

In the case of 1D systems, disorder is only in the lengths of junctions, and there is no room for the change in the topology of the network. The edge lengths are ℓ_*i*_ = ℓ_0_ + *ϵ*_*i*_, where *ϵ*_*i* ∈_ [−*ϵ*_0_, + *ϵ*_0_] with *uniform* distribution, and *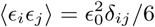*. In 2D the quenched disorder refers to (a) unequal edge lengths, and (b) topological defects defined as the local non-hexagonal polygons tiling the plane. The level of quenched disorder is controlled by randomizing the sites of a triangular lattice, based on which the polygonal lattice is generated using Voronoi tessellation. The edge lengths of the Voronoi lattice *ϵ*_*i*_, and density of defects *n*_*d*_ are then obtained by ensemble averaging over the ideally all the realizations of the disordered triangular lattice. Perturbing the sites of a triangular lattice, for the sites of the Delaunay lattice we have: **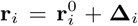**, with 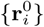. The spatial disorder term Δ_*i*_ is uniformly distributed in range [−Δ_0_, +Δ_0_], with local correlations: **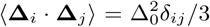**, where *δ*_*ij*_ is the Kronecker delta function. An ordered triangular lattice would return an ordered hexagonal lattice by Voronoi tessellation. By displacing randomly the sites of triangular lattice, we can distort the resultant Voronoi lattice. In order to obtain the disorder statistics of the Voronoi lattice, i.e. variations in the edge lengths *E*_*i*_, as well as the density of defects *n*_*d*_, we average over ensemble of disordered triangular lattices. Topological defects with finite (i.e. nonzero) density of defects in the thermodynamic limit, appear above a certain threshold of disorder Δ_*d*_ ≃ 0.25, in underlying Delaunay lattice (see Fig. (C.1)). The edge-length disorder in the Voronoi lattice increases linearly with Δ_0_ for Δ_0_ ≲0.5.

**Figure C.1:**
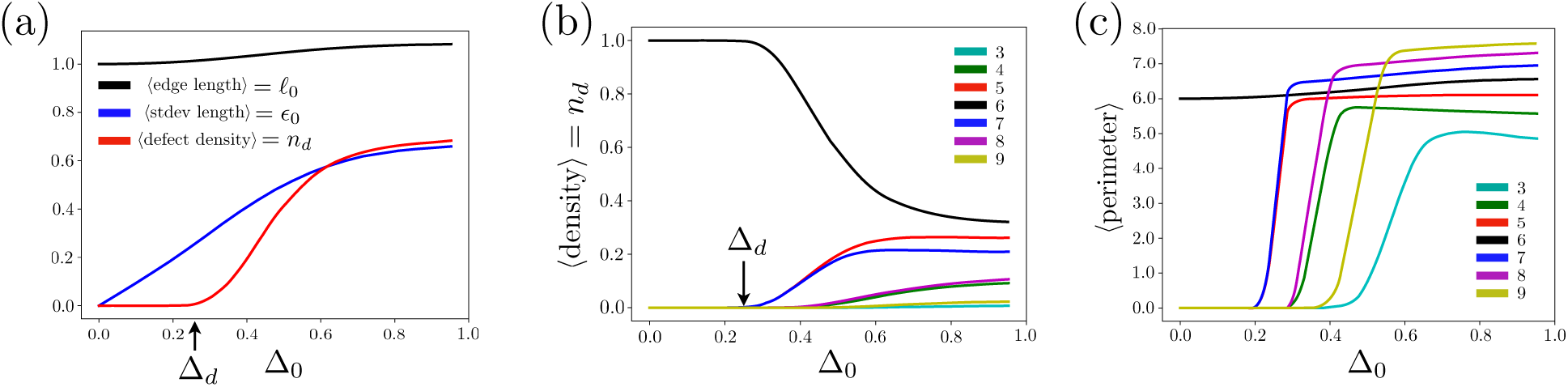
(a) Black curve represents the ensemble averaged lengths of edges versus the disorder in the position of cell centroids, This curve, as expected, is approximately one sixth of average perimeter of hexagons, depicted in (c), except for very large magnitudes where the effect of defects are become more important. Blue curve, shows *ϵ*_0_, the standard deviation (measure of disorder) of the edge length in the actual lattice. For small values of disorder Δ_0_ ≲ 0.5, the edge length disorder grows linearly, with the slope equal to one, and decreases for larger values of disorder. Red curve corresponds to the ensemble average total density of disorder as the disorder is enhanced. The disorders start at around Δ ≃ 0.25. (b), (c) Ensemble averaged densities and perimeters of polygons with different number of sides versus the disorder in the position of cell centroids. The ensemble average is carried out over 10000 realizations of 50 × 50 lattices.

### D Mean-field and Numerical Solutions

We define the mean-field approximation in this system as constant *a*^*f*^ *b*^*f*^ in space. As mentioned in the main text, the validity of this assumption follows from the diffusion-like dynamics of *p, s*. Bound proteins redistribute across the tissue until a relatively smooth state is reached, therefore the free proteins *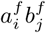* too, distribute uniformly. We also test out numerically, the validity of this assumption in 2D; see Fig. (D.2a). Here we first elaborate on the MF solutions in 1D, then discuss the 2D case where the MF solutions are divided into two different classes: trivial and nontrivial.

#### D.1 Mean-field solutions in one dimension

We start with one dimension for reasons that were discussed in main text. In one dimension the cells are juxtaposed in an array and are separated by junctions. The proteins localize on both sides of these junctions and form dimers. The lengths of the junctions are identical in ordered and random in disordered systems. A general scheme of one-dimensional arrays can be seen in Fig. (D.1). We use label *i* for cells, and the edge between the cells *i* and *i* + 1. Thus, ℓ_*i*_ = ℓ_0_ + *ϵ*_*i*_, with ℓ_0_ = 1 the unit of length, and *ϵ*_*i*_ the disorder. The reaction-diffusion (RD) equations governing *u* and *v* complexes on a junction *µ* shared by cells *i* and *j* read:

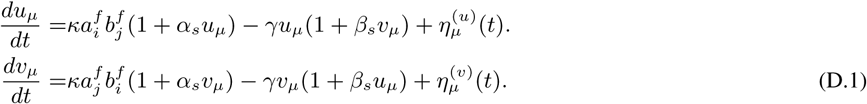

The above equations were originally proposed by Mani, et.al. in Ref. [38], namely our general RD equations, in ordered one-dimensional systems and in the limit of strictly local interactions, reduce to that in Ref. [38]. Here we briefly reproduce their results and move on to two dimensions. Starting with ordered systems, we first introduce the mean field (MF) solutions, in which the concentration of free A and B are uniform. Thus for MF solutions we drop the indices: *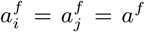* and similarly for *b*. Furthermore, in steady state, the concentrations of dimers are going to be uniform as well: *u*_*µ*_ = *u* and *v*_*µ*_ = *v*. Using the uniformity of, say *v*_*µ*_ = *v*_*µ±*1_, and that *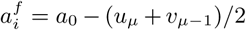*, one can write *a*^*f*^ and *b*^*f*^ in terms of *s* = *u* + *v*, namely: *a*^*f*^ = *a*_0_ − *s/*2, and *b*^*f*^ = *b*_0_ − *s/*2. Finally, using the definitions of *p, s*, we derive their dynamic equations:

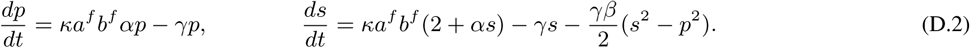

In the phase-space of the system, *p* − *s* plane, there exists a trivial class of solutions where *p* = 0. Plugging this in Eq. (D.2b) we get:

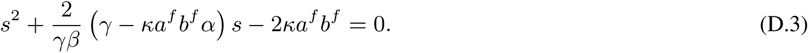

There also exist other branches of nullclines where *p* ≠0, thus *κa*^*f*^ *b*^*f*^ *α* = *γ* from Eq. (D.2a):

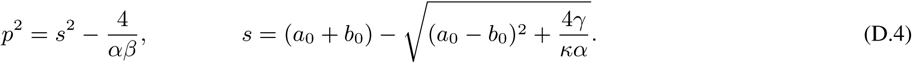

For *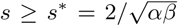*, the polarization is nonzero, and therefore, from Eq. (D.2a), we conclude that *kαa*^*f*^ *b*^*f*^ = *γ*, from which we derive Eq. (D.4b). The branches of solutions coincide at the bifurcation point, where the coefficient of the linear term in the last equation vanishes, i.e. *γ* = *κa*^*f*^ *b*^*f*^ *α*, corresponding to onset of polarization. Using the above relation we can determine the critical value in terms of *b*_0_.

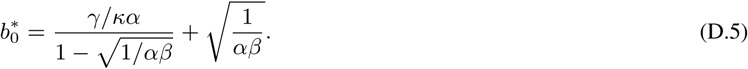

**Figure D.1:**
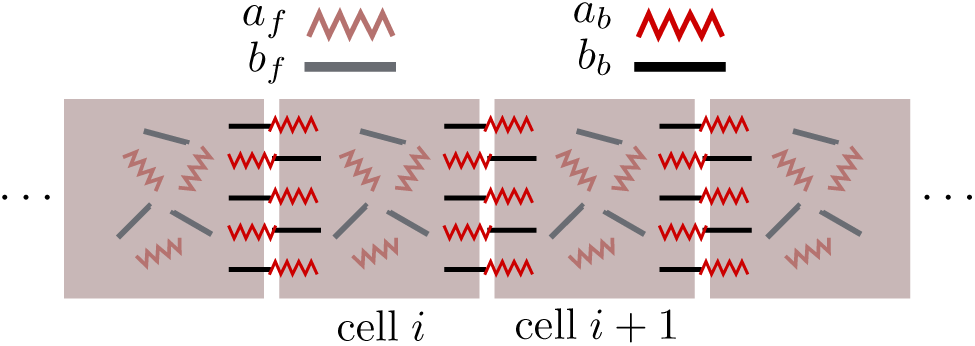
A cartoon of a one-dimensional array of cells. The free PCP proteins with concentrations *a*_*f*_, *b*_*f*_, are available to bind at the junctions. The PCP proteins bind at the interfaces. The sum of concentrations of [A-B] and [B-A] dimers at each junction is the total concentration *s*_*i*_ = *u*_*i,i*+1_ + *u*_*i*+1,*i*_, and the difference is defined as the polarization *p*_*i*_ = *u*_*i,i*+1_ − *u*_*i*+1,*i*_.

Note that in the above equations, for *αβ* = 0, the bifurcation point diverges.

#### D.2 Trivial mean-field solutions in two dimensions

Defining the mean-field approximation as the translational invariance of polarization along each of the three axes separately, the RD equations take the same form as in 1D, except the total amount of proteins A and B are shared by six edges instead of two. Therefore, as mentioned in the main text, the solutions too, resemble those in the 1D case. Each edge carries the same *p, s*, and the net polarity of each cell equals *p*_*c*_ = *p*_*e*_(1 − 2 cos *θ*), with *p*_*e*_ is the magnitude of polarization of one edge, calculated above, and *θ* is the angle between the two adjacent edges in ordered hexagons, which equals 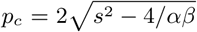, for *θ* = 2π/3. Here we consider a different situation in which edges no longer have identical *α, β*s. At this point, different values of *α*, and *β* can have various origins that are beyond the scope of our discussion. However in the Appendix Sec. (D.2), we argue that unequal parameters can be a consequence of, for instance, nonlocal interactions in elongated cells.

We assume the three pairs of parallel edges acquire coefficients *α*_1,2,3_ and *β*_1,2,3_. Without loss of generality we consider two scenarios: (i) *α*_1_ > *α*_2_ = *α*_3_, and (ii) *α*_1_ = *α* _2_ > *α*_3_. From the results we found in the case of sixfold symmetric lattices, the onset of bifurcation is inversely proportion to ^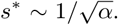^. Therefor in case (i), axis 1 is the first axis that shows instability upon increasing *b*_0_ above *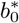*. Therefore, we have *a*^*f*^ *b*^*f*^ = *γ/*(*kα*_1_), where,

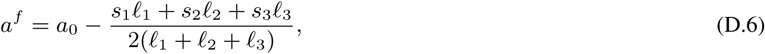

and similarly for *b*^*f*^. Again, as derived above, the axes where the polarization is zero, i.e. 2, 3, we have:

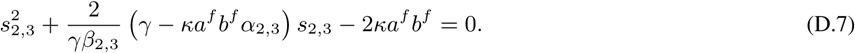

Using the fact that *ka*^*f*^ *b*^*f*^ = *γ/α*_1_, we get:

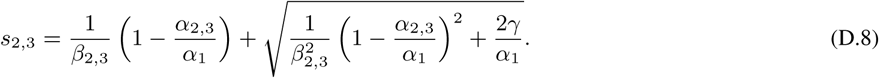

Defining *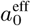* and *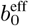*,

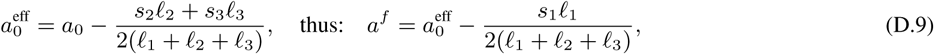

and similar expressions for *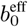* and *b*^*f*^, we get for *p*_1_, *s*_1_:

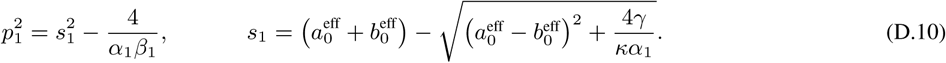

From the above analysis, we learn that when coefficients *α* and *β* of one pair of the edges are larger than those of the other two pairs, the system essentially reduces to a one dimensional problem, with effective pools of proteins A and B. There remains one more interesting case in which *α*_1_ = *α*_2_ > *α*_3_. In this case, the third axis remains unpolarized as the other two are effectively more absorbent, and share the total bound proteins, thus are equally polarized with four different degeneracies. As mentioned above, one situation of interest in which the coefficients *α, β* acquire unequal values for the edges is the elongated tissues. In such systems, the above two cases correspond to Figs. (E.1a) and (E.1b,c), respectively.

#### D.3 Nontrivial mean-field solutions in two dimensions

The nontrivial solutions in 2D do not possess the translational invariance of dipoles, and satisfy a weaker constraint. For reasons that become clear shortly, it is more insightful to use the junctional polarization in analyzing this case. As we will see, the full analytic solution to this problem is cumbersome. We only briefly touch upon this subject to provide some intuition into how large the basin of attraction is, for a system in SLI limit. The only assumption in nontrivial MF solutions is that *a*^*f*^ *b*^*f*^ is uniform across the tissue, which implies that the amounts of bound A and B are also equal across the tissue. This assumption is intuitively justified by the fact that linearized equations governing *u* and *v* obey diffusion-like equations in the continuum limit [38]. Therefore, the free proteins, too, spread diffusively into a rather uniform state. Furthermore, we numerically solved for *a*^*f*^ *b*^*f*^ in the steady state of systems with random initial condition. A generic distribution is illustrated in Fig. (D.2a). The quantity *a*^*f*^ *b*^*f*^ is almost uniformly distributed across the system. In SLI limit and/or for ordered lattices, by virtue of sixfold symmetry, all junctions are equally absorbing the proteins. Therefore above the bifurcation point, the net polarization *p* and the total amount of proteins *s*, of all junctions are identical. The only constraint is thus, three junctions have net positive polarizations and the other three have negative polarization; three outgoing and three incoming dipoles. In order to make it easier to picture such a configuration of dipoles, imagine we start from a trivial solution of type I, where three adjacent edges carry positive dipoles and the other three the negative ones. Now flipping one of the positive dipoles breaks the MF assumption of *a*^*f*^ *b*^*f*^ = constant. Therefore one of the negative dipoles must be flipped too. Since each dipole is shared between two cells, in order to satisfy the constraint in any finite-size system, this flipping process must continue until it forms a loop ending at the initial edge.

**Figure D.2:**
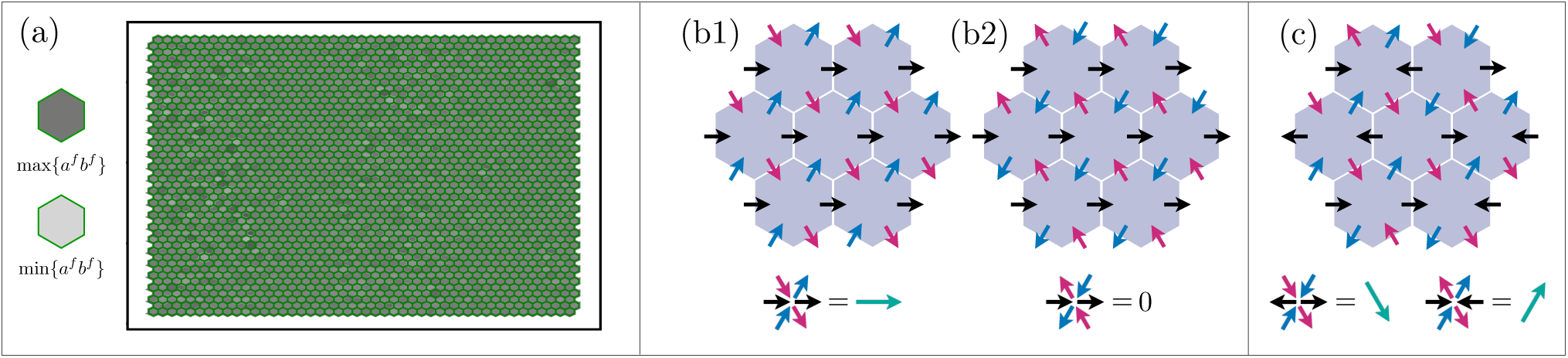
The left panel (a) shows the steady-state product of the concentrations *a*^*f*^ *b*^*f*^ for each cell, for a system with random initial state. The product remains almost uniform across the system, lending more support to the MF approximation. (b1) and (b2) illustrate the trivial MF solutions with nonzero and zero net polarizations respectively. (c) corresponds to a nontrivial MF solution. Each cell carries three incoming and three outgoing dipoles satisfying the MF assumption.

It is easy to see that the paths can be broken down into self-avoiding loops. Furthermore, all such loops preserve the net polarization of the tissue, set by the value of the control parameter *b*_0_*/a*_0_. However, they disrupts the uniform polarization of the trivial solutions, namely create excitations above the uniform configuration. Incidentally this is what we observe in simulations; the net polarization at the steady state is to a high accuracy independent of initial configuration. The small deviations is however understandable since the constraint of constant *a*^*f*^ *b*^*f*^ is not guaranteed to be accurately satisfied in real systems with arbitrary initial conditions. Moreover, in an ensemble of the systems starting from all possible initial conditions, all nontrivial configurations preserving the constraint as well as the net polarization, are equally accessible to the system.

The above arguments clarifiy why the system in SLI limit is not guaranteed to reach a state with large-scale polarization. Interestingly this simple analysis provides insight into why NLI mechanism stabilizes the states with nearly uniform polarization. Adding nonlocal interactions kills the nontrivial solutions of SLI case, by promoting the adjacent cells to have similar polarizations (segregation). Therefore, all the cells not only satisfy constant *a*^*f*^ *b*^*f*^, but also favor equal magnitudes of polarizations |**P**_*i*_.| In order for the cells to meet the two criteria simultaneously, the directions of the cell polarizations must also be parallel. In SLI case, however, different cells can have different |**P**_*i*_|, hence different orientations.

#### D.4 Unequal interaction ranges

We assumed throughout the main text that the range of nonlocal cytoplasmic interactions are identical for both intra- and interspecies kernels, i.e. *λ*_*uu*_ = *λ*_*uv*_. Here we consider cases in which one of the length scales is much smaller than the other: (a) *λ*_*uv*_ ≪ *λ*_*uu*_ = 0.5ℓ_0_, and (b) *λ*_*uu*_ ≪ *λ*_*uv*_ = 0.5ℓ_0_, namely one of the interactions falls in the SLI regime, while the other remains of NLI type.

We observe that when inhibitory interaction are short-ranged, i.e. (a), domains of correlated dipoles form of sizes roughly equal to 5-10 cell diameters, beyond which the correlation falls rapidly; Fig. (D.3a). Therefore, although the dipoles and clusters of aligned dipoles form, the coarsening stage of dynamics is not accomplished. In the opposite limit, the polarity becomes correlated on larger length scales, comparable to the case of *λ*_*uu*_ = *λ*_*uv*_, with minor modulations; Fig. (D.3b).

What we see in (a) is not unexpected. When the inhibitory interactions are local, the segregation is not successfully accomplished consistently over large distances, hence the small correlation length. On the contrary, (b) that roughly speaking, corresponds to a “local-activation, global-inhibition” (LAGI) mechanism, is expected to work properly. Indeed one would näively expect LAGI to work even better than equal interaction ranges. Nonetheless, we see that the latter indeed works better. The reason is as follows: The final state of a long-range polarized tissue, consists of cells that are effectively divided into two compartments carrying A and B separately. Now, suppose that most of A proteins are initially localized on the right side of a cell. Regardless of local or nonlocal interactions of like complexes, A and B push each other to the opposite sides of the cell using the nonlocal mutual inhibition. If this repulsive interaction is accompanied by a simultaneous attraction of A towards the initial locus of A (and the same for B), the segregation is facilitated and is more stable. Thus, nonlocal attraction (within a certain range) can amplify the segregation process.

Another important point to note is that, in both cases of *λ*_*uu*_*/*ℓ_0_ ≪ 1 and *λ*_*uu*_*/*ℓ_0_ ≃ 0.5, if the range of inhibitory interactions exceeds a certain length scale, roughly the size of a cell *λ*_*uv*_ ≳ ℓ_0_, the opposite sides of the cell inhibit each other, e.g. A on the right side, inhibits B on the left side. Thus, the B proteins sitting on the left side of the same cell are destabilized by A from across the cell.

**Figure D.3:**
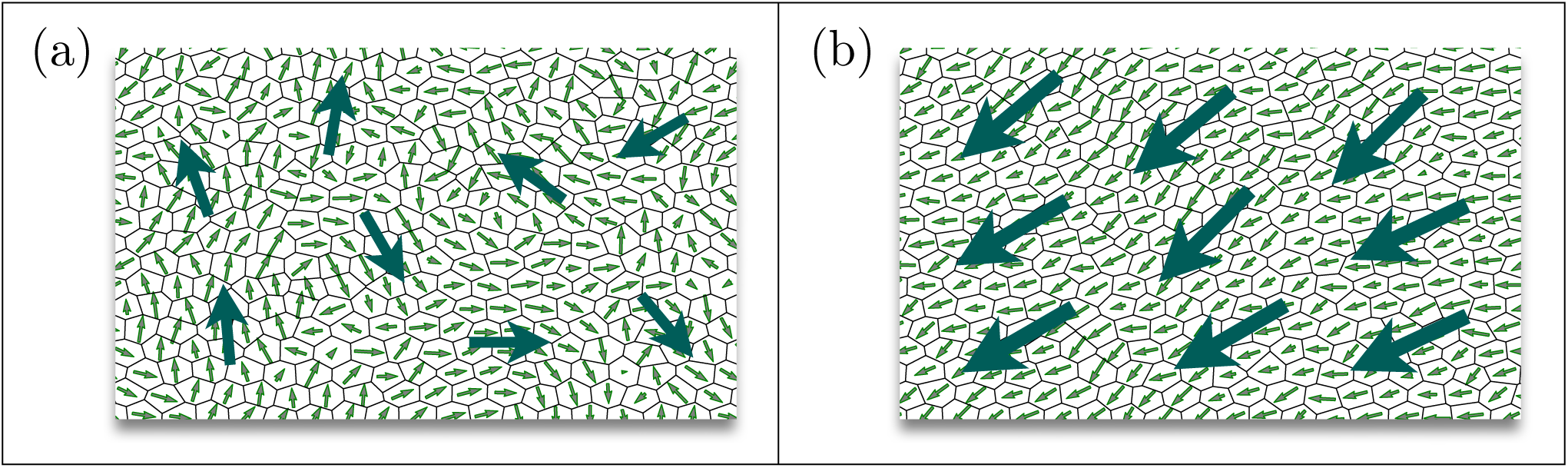
The steady-state solutions of polarization field in tissues where the ranges of nonlocal cytoplasmic interaction are unequal. (a) Inhibition between the unlike complexes are short-ranged (*λ*_*uv*_ */*ℓ_0_ = 0.01), while activation of the like complexes remains in NLI regime (*λ*_*uu*_*/*ℓ_0_ = 0.5), and (b) illustrates the exact opposite of (a). The big arrows represent local averages of polarization over regions of around 5 - 10 cell diameters.

However, as far as polarity is concerned, the B protein on the opposite side of the cell is indeed contributing positively to the same direction of polarity as A. Therefore, as mentioned in the main text, for *λ/*ℓ_0_ ≳ 1, the polarization, even if initially correlated over long distances, becomes highly unstable against any stochastic noise.

#### D.5 External cues

The external cues can appear as either bulk or boundary signals, each of which may be persistent or transient. Here we discuss and compare the responses of systems in SLI and NLI regimes, with different types of cues. In a nutshell, nonlocal cytoplasmic interactions (NLI) assist with the detection of global signals, even in highly disordered cases, whereas in SLI regime, only tissues with small disorder respond to global cues. Given that long-range polarizations have been observed in stages of development in which tissue is still highly disordered, NLI seems to be the necessary mechanism to detect the global cues.

##### Bulk signals

Bulk signals are incorporated in our model in the form of a constant gradient. In the absence of geometrical disorder of the tissue, the bulk cues are capable of rotating dipoles in both SLI and NLI regimes. However the NLI systems (a) require a much smaller strength of the signal, and (b) respond on a much shorter timescale than SLI systems. Upon increasing the geometrical irregularities, while the behavior of NLI systems remains the same with minor changes in the timescales, i.e. slower dynamics, the SLI fails to align with the gradient. Therefore, we conclude that, nonlocal cytoplasmic interactions play the key role in detecting the bulk cues, and aligning the collective polarization accordingly.

##### Boundary signals

In the SLI regime with small geometrical disorder, and in the absence of stochastic noise, we observe that a boundary signal creates a polarization wavefront that travels into the bulk. Consider a polarization wave as it travels to the right. As the wave-front approaches a column of cells, proteins of one type (say A) are absorbed to the left junctions shared with the cells in the previous column, and leave a net positive concentration of the other protein (B) on opposite side of the cell, which in turn attract A proteins to the left edges of next column and so on. Any finite value of stochastic noise, however, destroys the polarization beyond a length scale determined by the distance from the critical point.

The above results are true for persistent signals. When the signal is transient, the timescale of the signal becomes important. Suppose that the magnitude of the signal falls as an exponential *e*^−*t/τ*^*s*. In general, and as one would intuitively imagine, if *τ*_*s*_, is not much smaller than the dynamic timescale of NLI, found to be ∼50*/γ*, the signal manages to rotate the dipoles. Otherwise, the rotation is not fully accomplished. It goes without saying that SLI systems do not respond to transient signal.

### E Elongated Cell Geometry

In the case of anisotropic cells, as expected from the earlier discussions on the junctional coarse-graining, the renormalized value of different junctions are going to be dependent on their lengths. As in the case of one dimension, searching for mean-field (MF) solutions, we assume that in steady state, the concentrations of bound proteins are translationally invariant along the three main axes of the lattice. For cells with different lengths and thus different *α, β*’s, the polarization *p* in first equation of Eqs. (D.2), cannot vanish for all three different *α*’s. Therefore, the polarization along two of the axes remains zero, while the third (the longest) axis is polarized. Below, we investigate different possible scenarios for the two types of elongated cells, depicted in Fig. (E.1): the sixfold symmetry breaks down into twofold if elongation axis passes through a vertex (Fig. (E.1a)), or fourfold symmetries if elongation is perpendicular to two parallel edges (Figs. (E.1b, E.1c)). In both cases, the system behaves qualitatively like a 1D problem, and the polarization is perpendicular to elongation axis, pointing perpendicular to an edge (twofold) or towards a vertex (fourfold).

Previously we discussed the consequences of unequal *α, β*’s. The formalism and solutions are not directly applicable to the case of elongated systems with NLI, the reason being that the junction-junction interactions were not included in what we calculated in Appendix Sec. (D.2). Here, without deriving explicit expressions for the effective parameters in elongated system, we only argue that should we solve the full NLI equations in the elongated case, the longer junctions will acquire larger coefficients *α, β*. Note that in this self-consistent approach, the effective *α, β* are dependent on the concentrations of dimers on other edges. Therefore, assuming the system has reached its steady state, we can write the cooperative interactions as functions of the concentrations of dimers on all the edges, weighted by the geometrical factors originating from nonlocal interactions. For small ranges of NLI, *λ/*ℓ_0_ ≪ 1, the effect of other edges are negligible and only the self-interaction of each edge is to be taken into account. Intuitively and also from the expression given in Sec. (1), of the main text for nonlocal interactions, it can understood that the self-interaction is a monotonically increasing function of the edge length. Therefore longer edges with NLI, have larger values of *α, β*. Upon increasing *λ*, the mutual contributions between all pairs of edges increase. However the qualitative behavior of effective *α, β*’s for different edges does not change. With this in mind, one can see that the two situations discussed in Appendix Sec. (D.2), i.e. *α*_1_ > *α*_2_, *α*_3_ and *α*_1_ = *α*_2_ > *α*_3_ correspond to the cartoons in Figs. (E.1a), and (E.1b, E.1c), respectively. In (a) the polarization points toward the middle of the junction parallel to the elongation axis, whereas (b) polarization vector passes through a vertex. This analysis might explain, to some extent, the experimental observation of PCP vector pointing at a vertex. At any rate, cases (a) and (b) both acquire polarizations perpendicular to the elongation axis. There also exists an unstable configuration (c) which is unpolarized. The instability is again due to nonlocal interactions which forbids adjacent cells carrying opposite dimers; like the twofold degenerate trivial MF solutions of equilateral cells.

**Figure E.1:**
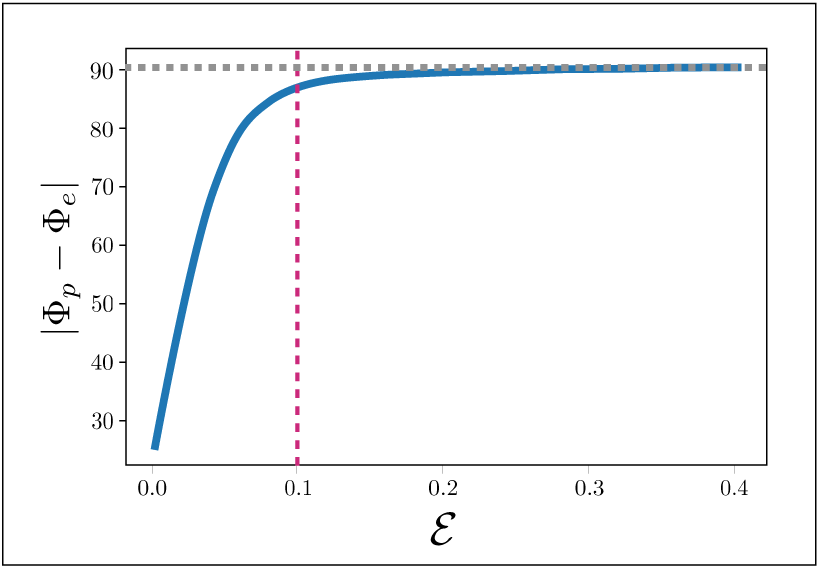
The angle between the axis of net polarization with *y*-axis, i.e. the axis of elongation, as a function of elongation index *E*, for a system with the same initial condition and primary lattice (see the explanation below, on the precise definition of ϕ_*p*_). At *ε* ≃ 0.1, the axes of polarization and elongation are almost orthogonal, with |ϕ_*p*_ − ϕ_*e*_| ≃ 87 (degrees).

#### E.1 Measure of elongation: cell nematic tensor

We discussed in the main text, how the emergence of perpendicular polarization can be explained in terms of nonlocal interactions of bound proteins. The cellular polarization as defined in Eq. (B.1), involves integration around the cell which is naturally dominated by the longer junctions, for uniform distribution of bound proteins. This is what we call a trivial shape effect. What is observed in experiments though, suggests perpendicular polarization beyond this trivial effect. The perpendicular polarization is not only a result of shape anisotropy, but indeed a consequence of larger absorbing power of long junctions. This can in principle be examined by comparing the density of dimers on long and short junctions. In practice, the best way to account for the shape anisotropy and discern its effect from that induced by NLI, is to calculate a nematic shape tensor (moment of inertia) of the cell, parametrizing the shape anisotropy of the cell.

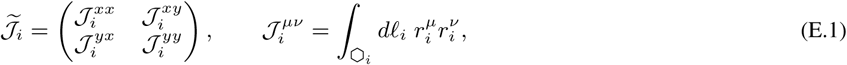

where *r* is measured with respect to geometrical center of mass, and its components *r*^*µ*^ and *r*^*ν*^ can each be *x* or *y*. The integration is carried out on the periphery of cell *i*, and *d*ℓ is the differential element of length, such that for constant radius we have *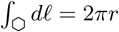*. In order to have the elongation index dimensionless, we normalize this tensor to the trace of *J*_*i*_, which is invariant under rotation. Thus, we have: *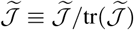*. One can find the elongation index and the angle with respect to *x*-axis, using the following relations:

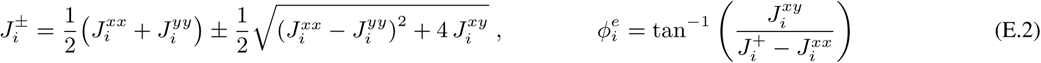

Elongation index is then defined as *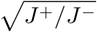*.

In order to compare our results to the experiments, we use the commonly used alternative representation of the elongation index, which is obtained by first making the *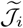* matrix, traceless:

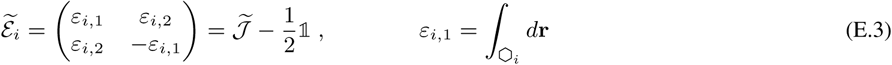

This is very much like what we introduced in Eq. (B.5). From the above equation, the magnitude of elongation and its angle to *x*-axis read,

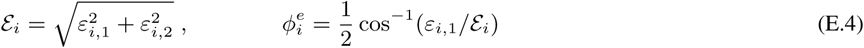

The elongation index is the spatial average of *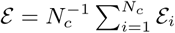*. With, *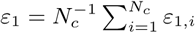*, the angle of nematic is defined as:

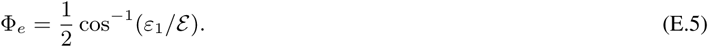

In terms of the elongation *ε* index, we plot in Fig. (E.1), the angle between *y*-axis and the steady-state polarization vector |Φ_*p*_ − Φ_*e*_ | (in degrees), versus different values of *ε*, for a system with the same initial condition and same “primary” lattice (i.e. before stretching), but elongated along *y*-axis. Here, Φ_*p*_ is defined as the angle of the axis between polarization and the *x*-axis, and over the domain of Φ_*p*_ ∈ [0, *π*). Since Φ_*e*_ = *π/*2 per definition, we have |Φ_*p*_ − Φ_*e*_|∈ [0, *π/*2]. Of course for *ε* = 0, the axis of elongation is not defined, yet we measure it with respect to *y*-axis. Furthermore, this figure is only an example in which the polarization for *ε* = 0, happens to make a small angle with *y*-axis, hence the pronounced effect of elongation. It is clear that depending on the topology of the lattice and initial condition, the polarization can be almost perpendicular to *y*-axis, even without elongation. Therefore among different simulations, we chose one with a relatively large effect of elongation.

It is noteworthy that for elongated cells with SLI too, the perpendicular polarization could be achieved due to the trivial shape effect; long edges cover wider angles, hence dominating the perpendicular component of the cellular polarization. However, in SLI case the proteins are by no means guaranteed to sit consistently on the long junctions; only on average they happen to be more localized on the long junctions.

### F Mutant Types and Their Corresponding Phenotypes

In order to further evaluate the validity of our proposed mechanism of signaling, we discuss, in this section, three other mutants, i.e. types III, IV, and V, and interpret their phenotypes in terms of their biological analogues. We demonstrate that our model reproduces the generic phenotypes of the corresponding mutants.

#### F.1 Type-III mutants: lack of cytoplasmic proteins

Type III suffers from lack of cytoplasmic proteins which is reflected as the suppression of the coefficients *α, β*, as both are ∞*c*_0_. As can be seen below in Fig. (F.1a), type-III phenotype shows mild non-autonomy. This mutant can be considered similar to *Dsh*^−^. It is very important to distinguish between types II and III, both of which involve cytoplasmic proteins. In type II, the nonlocal cytoplasmic interactions are suppressed, yet the membrane-bound complexes are able to interaction locally through SLI mechanism. Thus we interpret type II as a loss of function mutant. Type III, on the other hand, is depleted of cytoplasmic proteins and even local interactions are highly suppressed. Their phenotypes can be told apart by noting that the unlike type II, individual cells in type III are unpolarized, indeed similar to type I mutants. However both type I and II show very small non-autonomy, which is consistent with experiments. For images of phenotypes, see e.g. Ref. [35].

#### F.2 Type-IV mutants: clones with severed geometry

In type IV, a group of cells suffer from geometrical irregularities in the form of, (a) disordered lengths of junctions, i.e. (*ϵ*_0_), and size of the cells (apical area). In experiments [24], the irregularities decrease gradually as the distance from the center of the clone increases. Thus the clones do not define a clear boundary with the WT cells. In our simulations the level of irregularities drops according to a Gaussian function with the lengthscale of about 10 cells, around a nidus that marks the most geometrically disordered point. Experiments by Ma, et.al. [24], show that, in the absence of external cue (*ft*^−^, in *Drosophila* wing), the geometrical disorder disrupts polarization field by inducing a swirly pattern around the most severed region; see Fig. (1) in Ref. [24]. In Fig. (F.1b), we see that our simulations exhibit roughly similar patterns of polarity in the vicinity of the geometrically irregular cells.

#### F.3 Type-V mutants: double-mutants of membrane proteins

According to some experiments, e.g. [16, 42], the phenotypes of double-mutants in which both membrane proteins A and B are lacking, exhibit less non-autonomy that the single-mutants. This finding and its implications have provided important information regarding the mechanisms of intercellular signaling. The commonly accepted picture of the core PCP pathway has been the following: Vang acts as the ligand of Fz, and the direct signaling is monodirectional from Fz to Vang, namely in order for Vang to be localized at a junction, it must “sense” the density and “availability” of Fz in the adjacent cell, whereas Fz is oblivious of Vang concentration on the other side of the junction. The single mutants show non-autonomous phenotypes, which is also predicted by our model in type-I mutants. This picture has recently changed by comparing the phenotypes of single mutants *fz*^−^ and *Vang*^−^, with those of the double mutants *Vang*^−^*fz*^−^. The non-autonomy in the latter is greatly suppressed compared to single mutants. In a recent study [18], Fisher, et.al. have tested difference scenarios of Fz-Vang interactions, and concluded that no model based on monodirectional interactions can account for the suppression of non-autonomy in double mutants, and that in contradiction to the old picture, bidirectional interactions are necessary for the corresponding phenotypes to appear.

In light of the above-mentioned study and for reducing the complexities, originating from unequal signaling in the two direction, we constructed our simplified model originally based on the symmetric interactions between A and B. To check the validity of these assumptions, we examine our model’s prediction of double-mutants, by suppressing the concentration of the membrane proteins A and B in a clone. In agreement with experiments, we observe that the non-autonomous disruption of polarity is largely suppressed compared to single mutants, i.e. type I. This can be clearly seen in Fig. (F.1c), where the crescent of vanishing polarization in type I, is disappeared, and the polarization of WT cells is only slightly reoriented, wrapping around the clone.

**Figure F.1:**
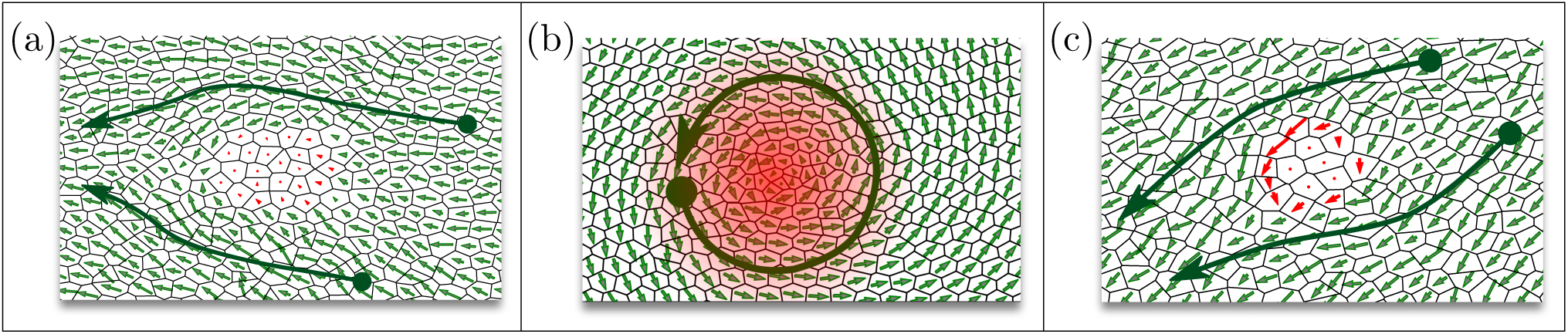
Illustrations of type III, IV, and V mutants in (a), (b), and (c), respectively. In all panels, the red regions represent the mutant cells. (a) Type III shows a clone with smaller values of *α*_mut._ = 0.01*α* and *β*_mut._ = 0.01*β*, which is interpreted as lack of cytoplasmic proteins, or their impaired modification by A and B. (b) shows the geometrically mutants that are centered around a highly disordered region. The resulting phenotype is a swirly pattern of polarization. (c) Double-mutants of both membrane proteins A and B. The non-autonomy is almost removed compared to to single-mutants type I.

#### F.4 Topological defects and domain walls in polarization field

Swirls (whorls) and saddles (crosses) are among repeatedly observed patterns in mutants [10, 24, 45–47]. In certain situations the polarization pattern contains topological defects like vortices, saddles, as well as domain walls separating two regions with different polarities. These patterns can arise either as transient or as long-lived states. The life-time of the defects in principle decreases with the magnitude of stochastic noise. As such the term “long-lived” here is referred to as *t* ≳ 1000*/γ*.

Starting from random initial conditions, systems with NLI are prone to formation of transient point defects that are observed at the interface of the coarsening domains of different polarizations; they subsequently disappear as the domains interact and align their polarities. In early-time dynamics, the dipoles interact mostly with dipoles in their respective longitudinal direction, because they point towards the edges carrying more dimers, and naturally interact more strongly with their neighbors positioned longitudinally. On the other hand, since dipoles are initially randomly oriented, the dynamics begin by forming small domains of parallel dipoles in all directions. Such point defects are thus a consequence of initial random orientation and stronger longitudinal correlations in early dynamics; see Fig. (F.2a). The appearance of transient defects in the polarization field was also predicted by Burak and Shraiman in [36].

The long-lived patterns might be the signature of a globally lacking ingredient, either insufficient B protein, *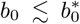*, or small diffusion length of the interaction-mediating proteins, *λ/*ℓ_0_ ≲ 0.1. In these cases, the defects form swirly patterns; Fig. (F.2b). These can be thought of as globally mutant systems. Line defects can appear as either line segments or closed loops. The line segments are bounded by two topological defects of charge 1*/*2. The two sides of the line carry opposite dipoles which are connected through U-turns at both ends. For instance, interesting patterns of polarization appear when the control parameter is tuned around the SLI critical point 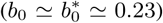. While in nearly ordered lattices, both cellular and global polarities remain close to zero, disordered lattices exhibit swirly robust patterns of polarizations, that are determined by the microscopic, i.e. cellular, geometry; see Fig. (F.2b). This is evident in patterns of polarization for a specific realization of quenched disorder, while *b*_0_ is changed in the vicinity of *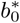*, i.e. 0.2 ≲*b*_0_ ≲ 0.25 (graphs not shown). The magnitudes of dipoles, however, grow as *b*_0_ increases. The swirls in this case are stabilized by the quenched disorders in the system. For larger *b*_0_, the long-range polarization arises smoothly as the nucleated domains grow, interact and align with one another.

Another type of long-lived defects that occur mostly in elongated systems with small disorder are domain walls separating two polarized regions, each with a net polarization perpendicular to the elongation axis. The reason why domain walls appear more easily in elongated systems is that elongation drives the system towards one-dimensional layers of cells that are less strongly coupled to cells in other layers than in isotropic systems; i.e. suppressing the effective coordination number of each cell. Therefore the domains with opposite polarizations can coexist. The occurrence and life time of the domain walls depend on the initial condition, as well as the level of the quenched and stochastic disorders. While in the presence of large quenched disorder, the dipoles of the dominant polarization conspire to swallow the opposite region, the latter may remain intact in ordered systems for long times until the stochastic noise removes it. Geometrical (quenched) disorder in each sample of finite size, serves as an infinitesimal bias (in addition to elongation), to break the +/− symmetry and remove the region of opposing polarities; Figs. (F.2c), (F.2d).

### G Methods

#### Simulations

Dynamical simulations are carried out using the forward Runge-Kutta method of 4th order. For each cell, starting from (typically) a random initial distribution of A and B proteins, we evolve the system according to the RD equations. All points on the edges of a given cell interact with each other through the kernels introduced in Sec. (2) of the main text. However since we assume the proteins are distributed uniformly along the junctions, it suffices to compute, for each realization of the lattice, the geometrical coefficients (𝒦_*µν*_) of junction-junction interactions by integrating the kernels along the two junctions, and for all pairs of junctions within a cell. Therefore, the integrations reduce to matrix products. Boundary conditions, in the systems without external cues of both kinds, are chosen to be periodic along both axes. The challenge arising from misalignment of boundary cells with disordered geometry (also pointed out in Ref. [24]), is circumvented using Voronoi tessellation, which generates cell layouts based on the seeded centroids. In systems with global (bulk) cues, we used free boundary conditions. In the case of boundary signal, the corresponding boundary is fixed while the others are free.

**Figure F.2:**
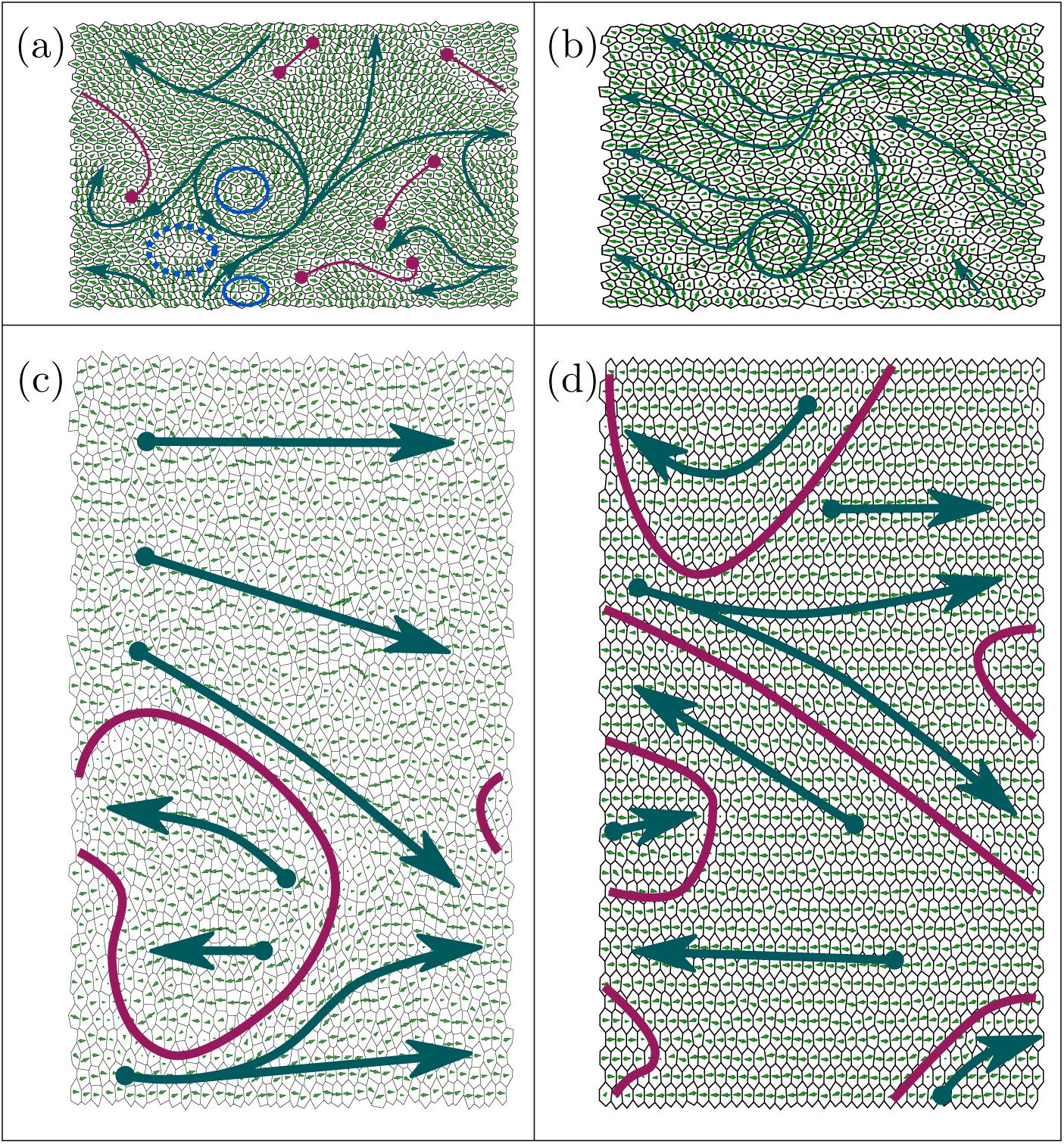
Illustrations of point and line defects in four different cases of systems with NLI of range *λ/*ℓ_0_ = 0.5. Green arrows show polarization flow fields. (a) An isotropic system during the early evolution (*γt* = 10), quenched disorder *ϵ*_0_ ≃0.5 and *n*_*d*_ ≃0.5. The solid-line circles represent vortices of topological charge 1, and the dashed circle has the charge −1. The red line segments with bullets at their ends are the line defects. (b) The steady state of a tissue with NLI, quenched disorder *ϵ*_0_ ≃0.5 and *n*_*d*_ ≃0.5, with low *b*_0_ = 0.35. This is another example of what was shown in main text Fig. (4b2), but at a larger value of *b*_0_, and above the SLI critical point. The net polarization is nonzero and pointing in this case to the left, yet defects like swirls are detectable. (c) and (d) show elongated systems ⟨ε⟩= 0.4. (c) Transient state (*γt* = 20) of a highly disordered system *ϵ*_0_ ≃0.5, and *n*_*d*_ = 0.5. Given the periodic boundary conditions in along both axes, one can see that the red solid curves form loops, i.e. domain walls, dividing the surface of a torus into two regions with opposite polarities. (d) The steady state of an elongated tissue with small quenched disorder of *ϵ*_0_ ≃ 0.18, and *n*_*d*_ = 0. The green arrows again represent the flow field.

#### Polarization and Correlation Functions

The cellular dipole moment is d<u>e</u>fined as the asymmetric angular distribution of A proteins inside a cell. The average of magnitudes *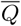* and the magnitude of averages *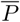*, are then calculated by spatial averaging over the lattice. Correlation functions and the corresponding correlation lengths are calculated using the definitions in the main text, Sec. (2). Due to the periodic boundary conditions, the correlation length is only defined up to the half the size of the lattice. We briefly mentioned in the text that during the early evolution of the polarization field, the longitudinal and transverse fluctuations have different correlation lengths. However, we have not separated the two, and have lumped them into one effective correlation length. As such, the spatial averaging over dipole-dipole correlations are only dependent on their radial distance.

